# Heat stress induces unreduced male gamete formation by targeting meiocyte translation

**DOI:** 10.1101/2022.08.11.503651

**Authors:** Cédric Schindfessel, Albert Cairo, Pavlína Mikulková, Chunlian Jin, Laura Lamelas Penas, Philip A. Wigge, Karel Říha, Danny Geelen

**Author notes:** Current affiliation: National Engineering Research Center for Ornamental Horticulture, Key Laboratory for Flower Breeding of Yunnan Province, Floriculture Research Institute, Yunnan Academy of Agricultural Science, Kunming, China (Chunlian Jin). The author responsible for distribution of materials integral to the findings presented in this article in accordance with the policy described in the Instructions for Authors (https://academic.oup.com/plcell/pages/General-Instructions) is: Danny Geelen.

## Abstract

Heat stress promotes the formation of unreduced (2n) male gametes through meiotic restitution, a driving force of evolutionary polyploidisation. Here we report that the molecular mechanism underlying heat tolerance of the meiotic division program in *Arabidopsis thaliana* relies on sustained protein translation of cell cycle genes. By leveraging natural variation in the *Arabidopsis* population, we identified heat-sensitive and heat-tolerant alleles of TARDY ASYNCHRONOUS MEIOSIS/CYCLINA1;2 (TAM). We show that TAM associates with specialised biomolecular condensates in meiotic cells under high temperatures. Through a mechanism that involves THREE DIVISION MUTANT1 (TDM1), TAM is required to maintain the translation of key meiotic cell cycle genes, including its own, thus preventing premature meiotic exit under heat stress conditions. Boosting TAM translation in heat-sensitive accessions using complementary peptides is sufficient to rescue the heat-induced defects. We propose that this mechanism can play a role in polyploidisation events and plant evolution in the context of the ongoing global climate change.

## INTRODUCTION

Plant genomes feature the remnants of past whole genome duplications that contributed to gene and species evolution (Jiao et al. 2011). Sexual polyploidization, through the fusion of diploid instead of haploid gametes, has been reported for a wide variety of plant species and is considered the main route to polyploidy (Bretagnolle and Thompson 1995; Kreiner et al. 2017; De Storme and Geelen 2013). The reductional division of chromosome sets during meiosis is central to the production of haploid male and female gametes and ensures ploidy consistency across generations in sexually reproducing organisms (Mercier et al. 2015). Therefore, diploid (2n) gamete formation is considered mainly a consequence of faulty meiotic chromosome segregation (d’Erfurth et al. 2008; Li et al. 2010; De Storme and Geelen 2011) or a premature termination of meiosis after the first meiotic division (Bulankova et al. 2010; Cromer et al. 2012; Sofroni et al. 2020). High environmental temperature conditions during meiosis promote the production of 2n gametes by evoking meiotic defects that cause either the first or second meiotic division to be skipped in a process called meiotic restitution (De Storme and Geelen 2013). Heat-induced meiotic restitution (HIMR) is a widespread defect in male meiosis extensively reported in plants (Pécrix et al. 2011; Wang et al. 2017; Schindfessel et al. 2021, 2023). Taken together with the observation that the timing of whole genome duplication events in the plant lineage is correlated with periods of climate change (Van de Peer et al. 2021) it has been suggested that HIMR could be a driving force for plant evolution in unstable environments.

In the light of the current global increase in temperature it is of interest to decorticate what genetic factors determine meiotic restitution (Schindfessel and Geelen 2025). Unreduced male gametes in Arabidopsis are formed by mutants defective in the cell cycle progression regulators TARDY ASYNCHRONOUS MEIOSIS (TAM/CYCA1;2) (Magnard et al. 2001; Wang et al. 2004), OMISSION OF SECOND DIVISION1 (OSD1) (d’Erfurth et al. 2009), and multiple CYCLIN DEPENDENT KINASES (CDKs) (Sofroni et al. 2020; Zhu et al. 2020). Mutations in these genes deregulate the meiotic progression pathway, that also involves the ANAPHASE PROMOTING COMPLEX/CYCLOSOME (APC/C), leading to the formation of dyad microspores. The timely termination of meiosis on the other hand requires the evolutionarily conserved nonsense-mediated RNA decay (NMD) factor SUPPRESSOR WITH MORPHOGENETIC EFFECTS ON GENITALIA7 (SMG7) and the plant specific protein THREE-DIVISION MUTANT1 (TDM1) (Bulankova et al. 2010; d’Erfurth et al. 2010; Cromer et al. 2012; Cairo et al. 2022). In meiocytes, TDM1 and SMG7 are loaded into P-bodies (PBs) and stimulate meiotic exit through sequestration of the translation initiation complex elF4F (Cairo et al. 2022). Remarkably, the inability of the *tdm1-3* mutant to exit meiosis is rescued by cycloheximide, providing evidence for a central role of translation inhibition in terminating meiosis (Cairo et al. 2022). TAM is a male meiosis specific cyclin that acts together with CDKA;1 during meiosis I to phosphorylate TDM1 and to prevent premature meiotic exit (Cifuentes et al. 2016). Mutants of *tam* skip the second meiotic division and produce dyads and unreduced pollen under standard growth conditions (Magnard et al. 2001; Wang et al. 2004). This phenotype depends on TDM1, suggesting that it relies on a premature activation of TDM1 to exit meiosis (Bulankova et al. 2010; Cifuentes et al. 2016).

Alternatively, meiotic restitution results from defects in chromosome segregation as observed in mutants of JASON, PARALLEL SPINDLE1 (PS1), DUET/MALE MEIOCYTE DEATH1 (MMD1) and FORMIN14 (AFH14) that regulate, separately or in combination, the organisation and orientation of the meiotic spindles (d’Erfurth et al. 2008; Li et al. 2010; De Storme and Geelen 2011; Andreuzza et al. 2015; Brownfield et al. 2015). In these mutants, meiosis I proceeds into meiosis II and during the second meiotic division parallel or fused spindles form, re-joining the sister chromatids from the homologous chromosome pairs.

The molecular mechanism by which heat affects meiotic exit to induce HIMR has not been elucidated. Here we exploited the natural variation in HIMR sensitivity in a population of *Arabidopsis thaliana* to identify TAM as key HIMR regulator that relocates to cytoplasmic molecular condensates under heat stress conditions. We show that TAM is required for maintaining meiotic protein translation at high temperature, in order to prevent a premature termination of meiosis and the production of unreduced gametes and polyploid offspring.

## RESULTS

### Natural variation in heat-induced meiotic restitution in Arabidopsis

To investigate HIMR in a natural plant population, we exposed 172 *Arabidopsis thaliana* accessions from different climatological regions (Supp. Figure 1a) to 32°C for 24h and compared their tetrad-stage of male meiosis to a 20°C control. The formation of reduced meiotic tetrads or unreduced triads and dyads (Figure 1a) was recorded to determine the incidence of defects in male meiosis under both conditions. Across these accessions grown under control conditions (20°C), a total of 2 dyads and 1 triad were recorded among 44405 tetrad-stage configurations viewed. All other microspores appeared as tetrads, showing that male meiosis robustly results in 4 haploid microspores at the end of meiosis in a wide variety of *Arabidopsis* accessions. After heat treatment a total of 8174 dyads, 1411 triads and otherwise tetrads were recorded out of 84975 microspores. Across the accessions, only 33 produced 100% normal tetrads after 24h at 32°C, indicating that meiotic restitution as a response to changing environmental temperature is a widespread phenomenon in *Arabidopsis* (Figure 1b). Remarkably, the dyad production after heat varied largely between accessions, with average dyad production ranging from 0% to nearly 100% (Figure 1b), indicating that intra-species genetic variation is responsible for the variation in meiotic restitution rates.

**Figure 1.**
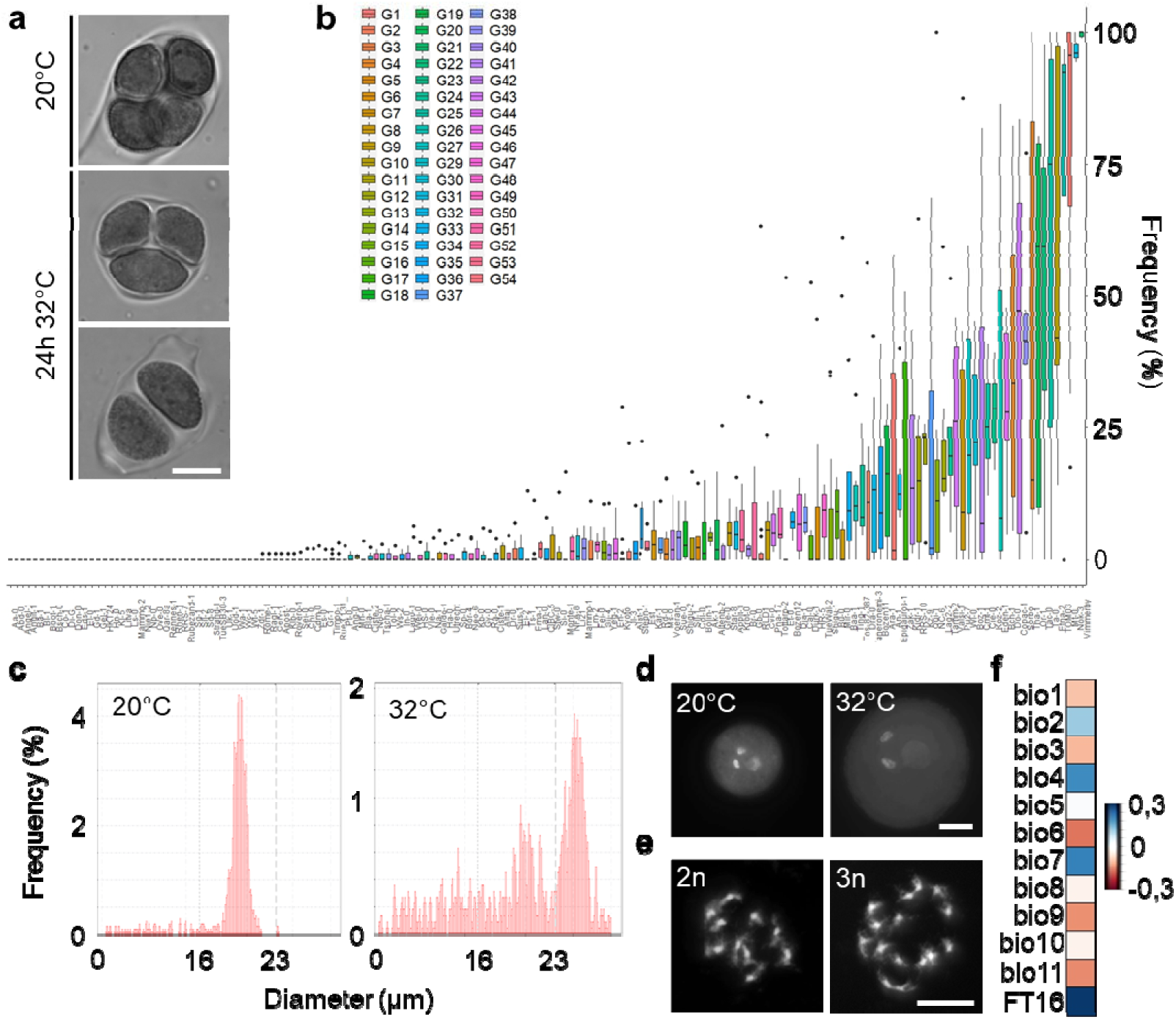
Natural variation in heat-induced meiotic restitution in Arabidopsis. **a)** Orcein stained tetrad-stage male meiocytes. Example of a tetrad under 20°C control conditions and a triad and a dyad after a 24h at 32°C heat treatment. **b)** Frequency of dyad production after 24h at 32°C for all 172 accessions analysed, ranked according to average dyad frequency. Boxes are coloured according to the batch the accession was analysed in (G1-G54). Lower and upper hinges of the box correspond to the first and third quartile. Whiskers extend from the hinges no further than 1.5 times the inter-quartile range (or to the highest or lowest data point). Data points beyond the range of the whiskers are plotted separately. **c)** representative example of the pollen particle size distribution histogram of Mt-0 plants kept at control conditions (20°C) or 8 days after a 24h at 32°C heat treatment **d)** DAPI stained images of control (20°C) pollen and enlarged pollen of Mt-0 after heat treatment. **e)** DAPI stained, enzyme digested mitotic chromosome spreads of a diploid (2n=10) and a triploid (3n=15) Mt-0 plant after heat treatment and self-fertilization. **f)** Spearman correlation heat map of dyad frequency at 32°C to 12 temperature-related climatological variables (bio1-11) and flowering time under 16h light (FT16). Details in Supp. Table 2. Count data used for (b) can be found in Supp. File 1. For (a, d, e) scale bar = 10µm.

In the most heat-sensitive accessions (Vimmerby, Nok-3, Mt-0, Tac-0, Ta-0, TOM03, Etna-2, Copac-1, Sorbo), we noticed the production of larger pollen grains in a brief time period around 7-9 days after the heat treatment (Figure 1c, d; Supp. Figure 1b-e, 2, 3). This timing as well as the presence of enlarged vegetative and sperm nuclei in these pollen suggest that the unreduced meiotic products we observed after the heat treatment developed further into mature 2n pollen (De Storme et al. 2013). The presence and viability of diploid pollen was confirmed for accession Mt-0 by using them in reciprocal manual crosses between heat treated and control plants as well as letting heat treated plants self fertilise. Triploid (3n) offspring were found only when the male parent or selfed plant had been heat treated (Figure 1e; Supp. Table 1) We extracted high resolution climatological data for all accessions from the CHELSEA Bioclim data set (Karger et al. (2017); Supp. Table 2) and data on flowering time from the Arabidopsis 1001 genomes project (Atwell et al. 2010). Dyad production at 32°C was positively correlated with flowering time and the annual and seasonal temperature range of the accession’s natural habitat. Negative correlations were found for parameters that measure mean temperature (Figure 1f; Supp. Table 2).

### Heat-sensitive TAM alleles lead to meiotic restitution

Using a bulked segregant analysis between accessions Ler (1% dyads at 32°C) and Mt-0 (80% dyads at 32°C) we mapped the locus responsible for Mt-0’s high dyad phenotype to TAM/CYCA1;2 (Supp. Figure 4; Supp. Table 3). The TAM defect in Mt-0 at high temperatures was confirmed by allelism test crosses with the *tam-1* and *tam-2* mutant alleles in the F1 (Figure 2a). Subsequently Mt-0 and Ler were used as tester lines to complement heat-sensitive accessions revealed in our initial tetrad-stage screen. For 7 accessions it was found that they were unable to complement the heat-induced dyad phenotype of Mt-0, whereas in crosses with Ler, the dyad phenotype was suppressed (Figure 2b). Finally, we cloned 5 natural TAM alleles (pTAM::TAM-GFP) from sensitive accessions (Mt-0, Nok-3, Vimmerby and Tac-0) and non-sensitive accession Col-0 and introduced them into the *tam-1* mutant in the Col-0 background. All alleles complemented the mutant at 20°C, but only the Col-0 allele was able to rescue the high dyad phenotype at 32°C (Figure 2c). These results validate TAM as the heat-sensitive gene leading to HIMR in at least 8 natural accessions of *Arabidopsis thaliana*.

**Figure 2.**
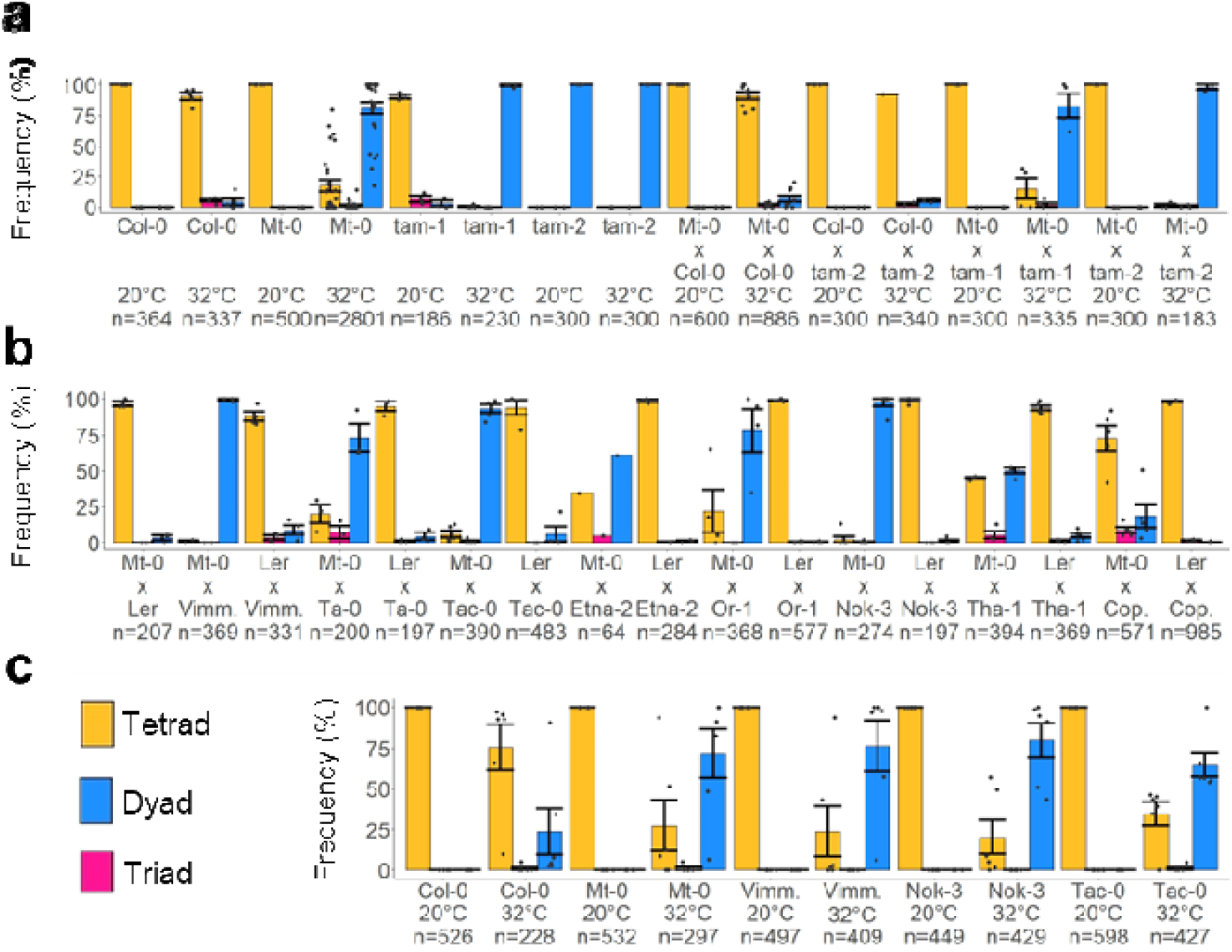
Heat-sensitive TAM alleles lead to meiotic restitution. **a-c)** Frequency of tetrad, triad and dyad formation at 20°C or after 24h at 32°C for various lines and F1 hybrids (a,b) or for tam-1 lines expressing natural alleles of pTAM::TAM-GFP of five Arabidopsis accessions (c). Data for Col-0 and Mt-0 in (a) where reused from Figure 1b. Error bars represent standard errors, n is the number of tetrad-stage configurations counted and the black dots represent individual flower buds.

### The TAM protein reacts to stress

To investigate TAM’s role at high temperatures we analysed TAM protein localisation using a Col-0 pTAM::TAM-GFP marker line in the *tam-1* background (Figure 2c, 3a). In both fixed tissue and live imaging TAM-GFP was present in the cytoplasm of leptotene meiotic cells reaching peak levels during mid prophase, followed by a gradual decline during late prophase (Figure 3a; Supp. Figure 5a). At the onset of the first meiotic division the TAM-GFP signal is cleared from the cells by proteasomal degradation (Cromer et al. (2012); Supp. Figure 5a; Supp. Movie 1). We noticed the appearance of faint cytoplasmic TAM-GFP foci during prophase, the visibility of which was enhanced in fixed samples compared to live imaging (Figure 3a). These foci became more prominent when plants were heat treated at 32°C or just before the meiotic cells began to die in live imaging, typically one hour before meiotic progression halted (Figure 3a). This suggest that TAM actively moves to these foci under stressful conditions.

**Figure 3.**
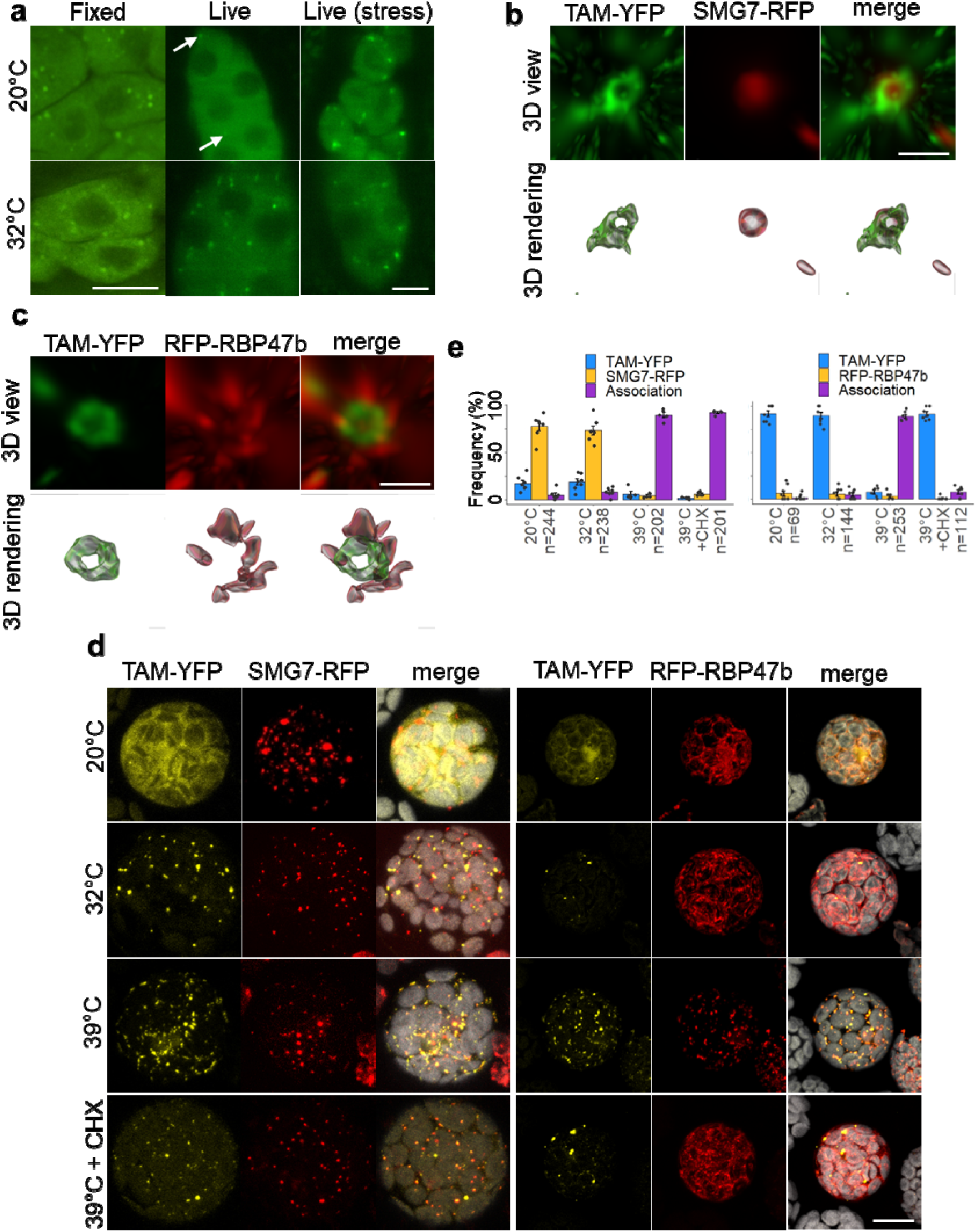
TAM associates with biomolecular condensate. **a)** Expression of pTAM::TAM-GFP of Col-0 in the tam-1 background after formaldehyde fixation or during live imaging using light sheet microscopy. Samples were fixed after 24h at 20°C/32°C, whereas live imaging snapshots are taken after 30min at 20°C/32°C. Stressed cells are samples that died in about 1h after the snapshot. **b-c)** Super resolution images and 3D rendering of the colocalization of pTAM::TAM-GFP and pSMG7::SMG7-tagRFP (b) or tagRFP-RBP47b (c) in prophase meiocytes. **d)** Expression and colocalization of p35S::TAM-YFP, p35S::SMG7-tagRFP and p35S::tagRFP-RBP47b in mesophyll protoplasts at various temperatures or treated with 100µM cycloheximide (CHX). Chloroplast autofluorescence indicated in grey. **e)** Quantification of the association described in (d). For (e) error bars represent standard errors, n is the number of individual foci counted and individual protoplasts (minimum 6) are represented by black dots. For (a, d) scale bar equals 10µm, for (b, c) scale bar equals 1µm.

**Figure 4.**
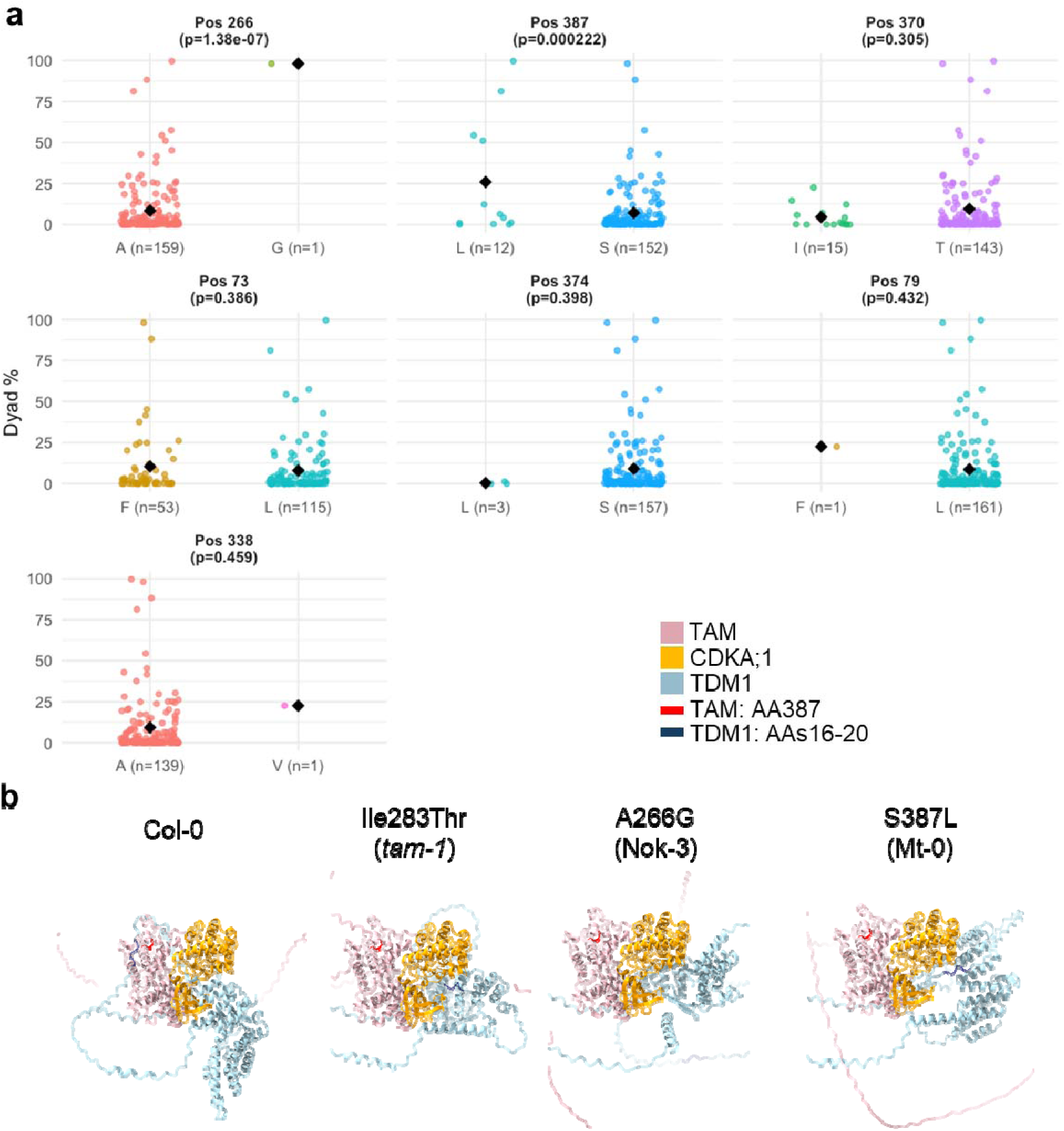
Heat-sensitive natural alleles have altered TAM protein confirmations. **a)** Comparison of Single Amino Acid Polymorphisms (SAAPs) with dyad frequency after 24h at 32°C (Supp. file 1) for every TAM SAAP in the 172 accessions tested (Sequence data from the 1001 genomes dataset (Alonso-Blanco et al., 2016) and our own sequencing data for Mt-0). p-values based on ANOVA comparing minor and major SAAPs, only SAAPs with p<0,5 are included in this figures, other SAAPs can be found in Supp. Fig. 8. **b)** Alphafold3 modelling of the TAM-CDKA;1-TDM1 complex for different TAM alleles. AA position 387 in TAM and the region around phosphorylation site Thr16 in TDM1 are indicated.

**Figure 5.**
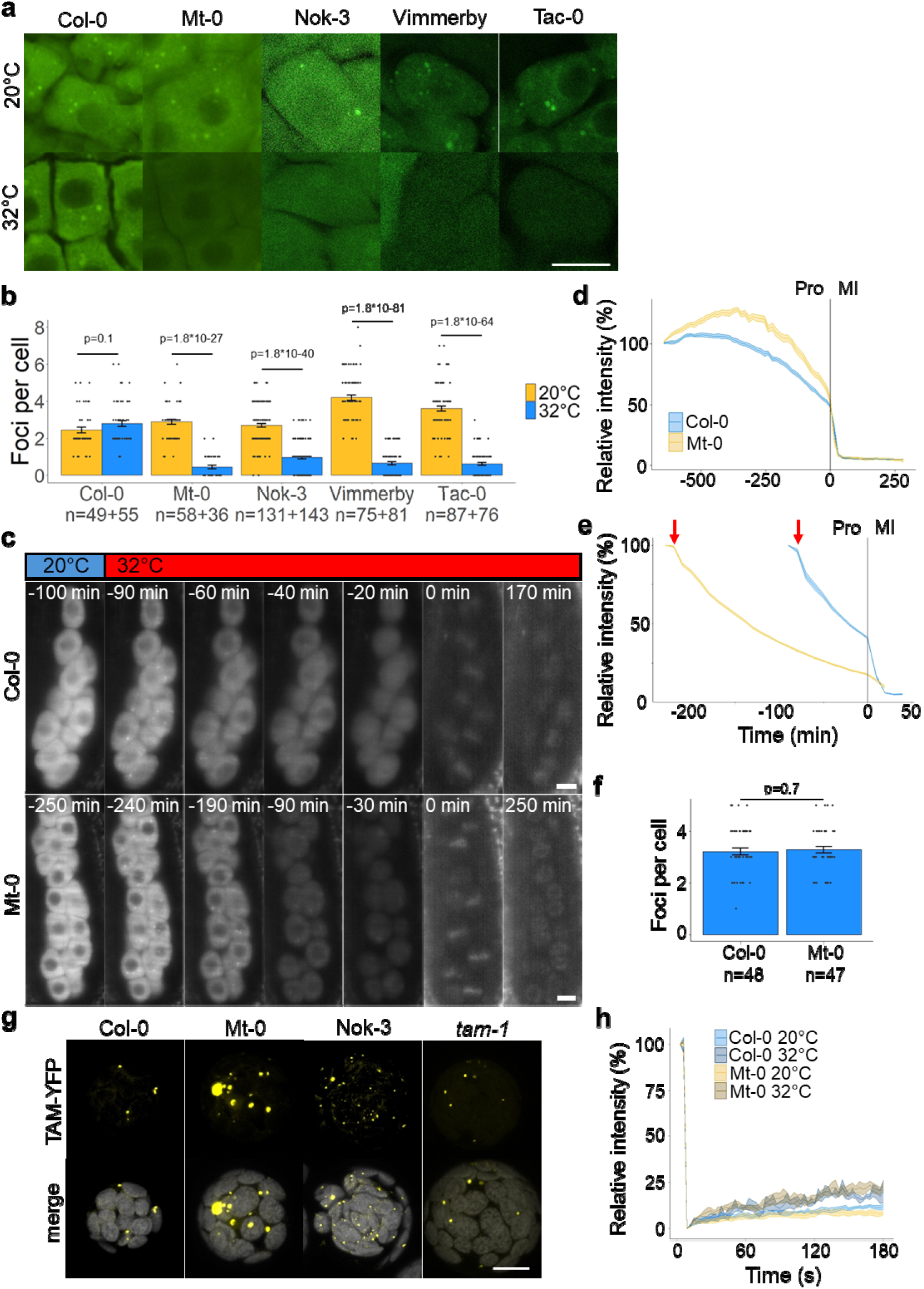
High temperature interferes with TAM protein expression. **a)** Expression of pTAM::TAM-GFP of five natural alleles (Col-0, Mt-0, Nok-3, Vimmerby, Tac-0) in prophase meiocytes in the tam-1 background at 20°C and after 24h at 32°C. **b)** Quantification of the number of foci per cell for (a). **c)** time-lapse of pTAM::TAM-GFP (Col-0 or Mt-0 allele) expressing meiocytes from late prophase to tetrad/dyad stage. Time point 0 denotes the entry into meiosis I. For the images of interkinesis and dyad/tetrad stage the brightness of the image was increased to allow visualisation of the autofluorescence in the organelle band. **d,e)** Quantification of the fluorescence intensity of TAM-GFP during the time-lapse experiment described in (c) for plants kept at 20°C (d) or during the temperature shift (e). Red arrows denote the shift from 20°C to 32°C. Pro = prophase; MI = meiosis I. The vertical line at time point 0 denotes the entry into meiosis I. **f)** Quantification of the number of TAM-GFP foci on time point 10min after the temperature shift as described for (c). **g)** Expression of p35S::TAM-YFP of three natural alleles (Col-0, Mt-0, Nok-3) and the tam-1 mutant allele in mesophyll protoplasts after 1h at 32°C treatment. **h)** FRAP analysis of TAM-YFP of Col-0 and Mt-0 in protoplasts after induction of TAM granules by 1h at 32°C treatment. The FRAP bleaching and recovery where either performed at 20°C or at 32°C using a heated microscopy slide. For (b, f) error bars represent standard errors, n is the number of meiocytes quantified and black dots represent individual meiocytes. p-values are based on Wilcoxon rank-sum tests, Benjamini-Hochberg adjusted for multiple testing. For (d,e,h) the error ribbon represents standard errors. For (d,e) n = 18-26 meiocytes from at least 3 different anthers, for (h) n = 3-5 foci from 3-5 different protoplasts. For (a, c, g) scale bar equals 10µm.

### TAM associates with biomolecular condensate in meiocytes

Multi-layered meiotic bodies (M-bodies; MBs; (Cairo et al. 2025)) in *Arabidopsis* meiotic cells are an association of two biomolecular condensates: a processing body core (PB; Chantarachot and Bailey-Serres (2018)) and a SG shell (Protter and Parker (2016)). To check whether the TAM foci are part of MBs, PB component SMG7-RFP and SG component RFP-RBP47b were stably co-expressed with TAM-GFP in meiotic cells. At both 20°C and 32°C TAM-GFP associates with SMG7-RFP, and RFP-RBP47b (Supp. Figure 5b-e) showing that TAM is a component of MBs. Super resolution microscopy on meiotic cells revealed that TAM-GFP localises in the periphery of SMG7-RFP at the core of the MB (Figure 3b). Curiously, super resolution colocalization with RFP-RBP47b showed that TAM-GFP is at the inside of the SG shell of the MB (Figure 3c). These results indicate that TAM resides at the interface between the PB core and SG shell of the MB.

To further dissect the stress-induced association of TAM with MB components we expressed p35S::TAM-YFP, p35S::SMG7-RFP and p35S::tagRFP-RBP47b in *Arabidopsis* mesophyll protoplasts, allowing for easier manipulation of temperature and pharmacological treatments to study condensate behaviour. At 20°C TAM was mainly cytoplasmic, forming foci in about 15% of the cells (Figure 3d, e; Supp. Figure 6a-c). These foci did not associate with SMG7 and at 20°C RBP47b does not form condensates (Figure 3d,e). At 32°C TAM formed condensates in 100% of the transfected cells (Supp. Figure 6b, c) and partially associated with SMG7 foci, but RBP47b did not form prominent condensates at this temperature (Figure 3d, e; Supp. Figure 6b, c). At 39°C TAM associated with SMG7- and with RBP47b-granules (Figure 3d, e). This shows that TAM associates with MB components in reaction to heat stress and indicates that TAM condensates form at lower temperatures than canonical SGs in somatic cells. To corroborate this observation we applied cycloheximide, an inhibitor of protein translation which prevents the nucleation of SGs by retaining the stalled mRNAs within polysomes (Supp. Figure 6b). Under these conditions TAM remained in condensed form and still associated with SMG7 at 39°C, whereas RBP47b-granules collapsed (Figure 3d, e; Supp. Figure 6b). These results where independently confirmed using the SG marker 35S::CFP-G3BP-2 (Supp. Figure 6d, e).

**Figure 6.**
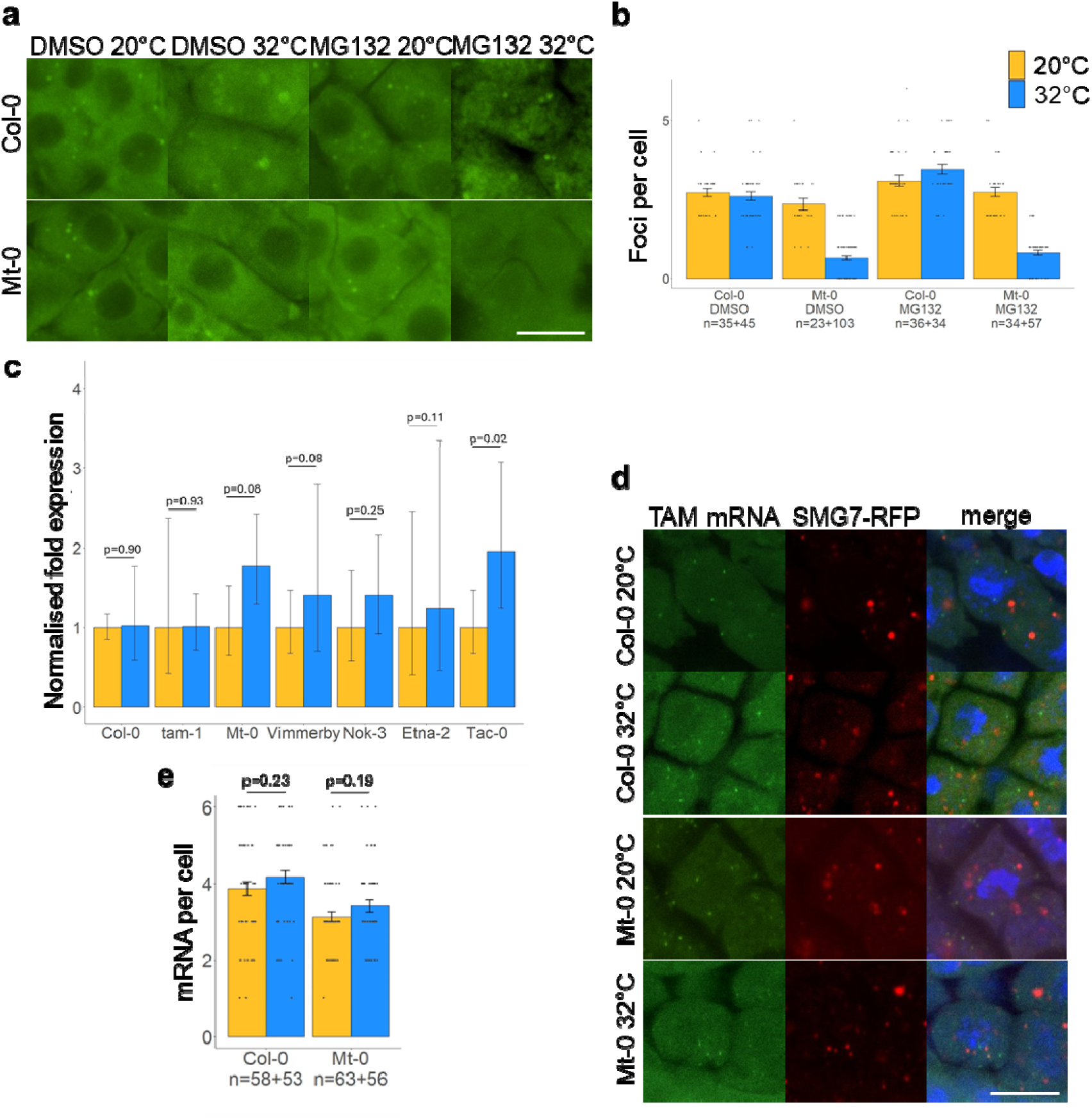
Analysis of TAM degradation and transcription. **a)** Expression of pTAM::TAM-GFP of Col-0 or Mt-0 in prophase meiocytes in the tam-1 background in vitro treated for 24h with DMSO or MG132 at 20°C or 32°C. **b)** Quantification of the number of TAM-GFP foci per cell for (a). **c)** qPCR-based relative expression levels of TAM in flowerbuds of natural accessions at 20°C or after 24h at 32°C; normalised to 20°C for every accession. **d)** Whole-mount RNA FISH labelling of TAM mRNA of Col-0 and Mt-0 at different temperatures in lines expressing pSMG7::SMG7-RFP, counterstained with DAPI. **e)** Quantification of the number of TAM mRNAs in (d). For (b,e) error bars represent standard errors, black dots represent individual cells and n equals the number of cells counted. p-values are based on (pairwise) Wilcoxon rank-sum tests, Benjamini-Hochberg adjusted for multiple testing. p-values for (b) are in Supp. Table 4. For (c) error bars represent standard errors, p-values correspond to a linear model fit comparing the 20°C control to the 32°C treatment, n=3 plants. For a,d scale bar equals 10µm.

Altogether, TAM localisation in somatic and meiotic cells shows that TAM translocates from the cytoplasm to molecular condensates in response to increasing temperature, independent from canonical SG formation. During meiosis, TAM associates with MBs and resides in the interface between the PB core and SG shell.

### Heat-sensitive natural alleles have altered TAM protein confirmations

It was recently demonstrated that the translocation of TAM to condensates is required for normal meiotic progression during heat stress and the formation of tetrad microspore (De Jaeger-Braet et al. 2025). The TAM protein contains an intrinsically disordered region (IDR) that was shown to be essential for this translocation. We therefore set out a comparative structural analysis of the temperature sensitive and resistant natural TAM alleles.

Sequence alignment of the 8 temperature sensitive TAM alleles revealed no common SNPs (Single Nucleotide Polymorphisms), compared to the Col-0 reference (Supp. Figure 7a). Furthermore, SNPs in the sensitive alleles are not unique within the Arabidopsis population and are often shared with more resistant accessions (Supp. Figure 7a-c). Therefore, a combination of SNPs is likely responsible for what constitutes a heat-sensitive TAM allele. To interrogate at the protein level how SAAPs (Single Amino Acid Polymorphisms) cause increased dyad formation, we extracted TAM protein sequences of the 172 accessions analysed (1001 genomes and our own sequencing data for Mt-0) and compared for each position SAAPs and average dyad frequencies (Figure 4a; Supp. Figure 8). Our results show two significant positions (t-test, p<0.05): Ala266Gly (p=1.38*10^-7^) present in Nok-3 and Ser387Leu (p=2.2*10^-4^) present in Mt-0, Vimmerby and Tac-0. These SAAPs occur in the N-or C-terminal cyclin fold domains of TAM (Supp. Figure 7c). Notably no significant SAAPs were detected in the IDR.

**Figure 7.**
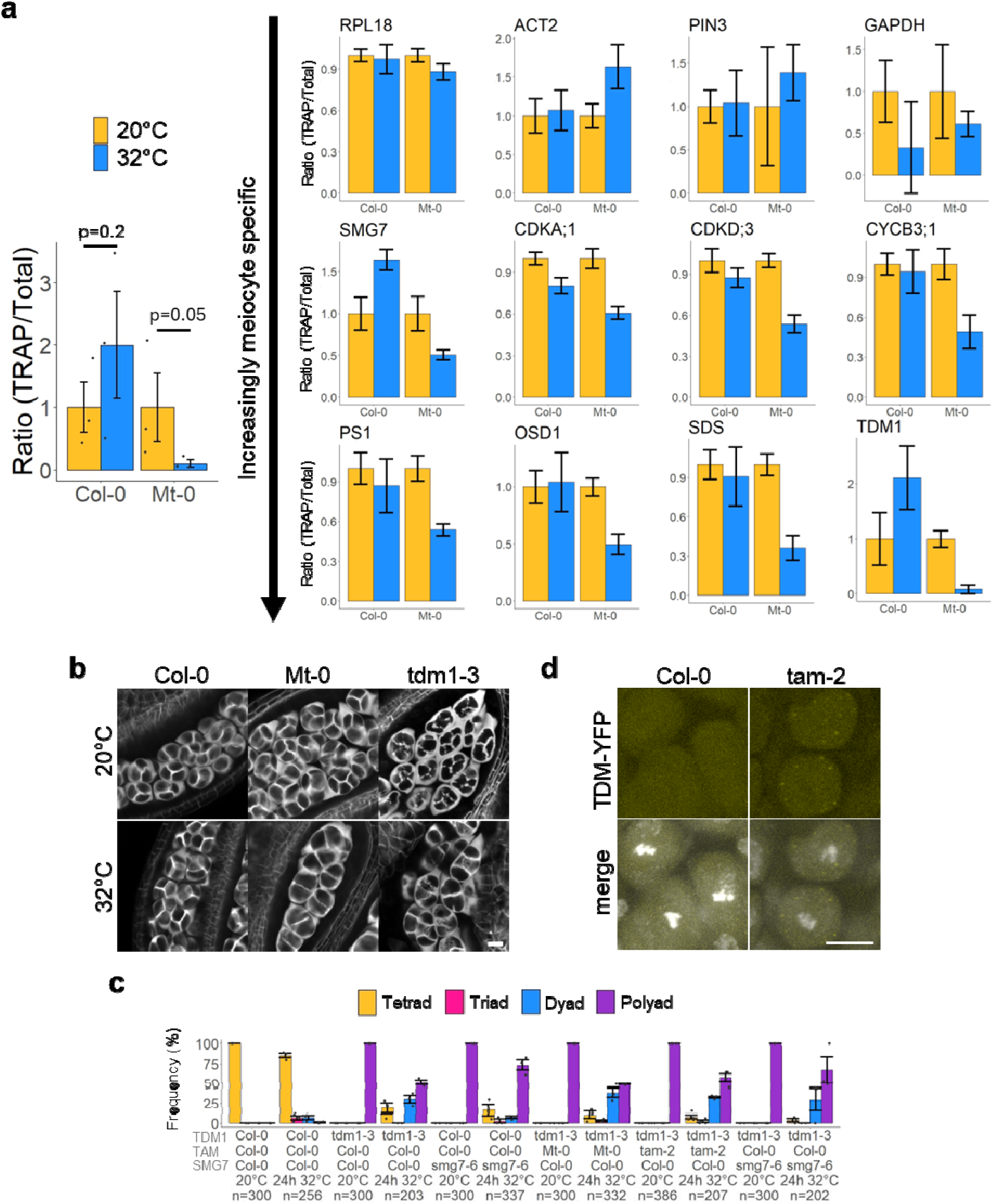
TAM is required to maintain meiotic translation at high temperature. **a)** qPCR-based ratio of the amount of TRAP vs total mRNA for various genes in Col-0 and Mt-0 lines expressing pUBQ10::GFP-RPL18; normalised to the 20°C control for every accession. Three independent biological replicates, each time pooling about 25 plants. Error bars represent standard errors, p-values are based on Wilcoxon rank-sum tests. **b)** SR2200 stained, tetrad-stage anthers of Col-0, Mt-0 and tdm1-3 at 20°C or after 24h at 32°C. **c)** Frequency of tetrad, triad, dyad and polyad formation at 20°C or after 24h at 32°C for lines with different homozygous combinations of alleles/mutations of TAM, TDM1 and SMG7. Error bars represent standard errors, n is the number of tetrad-stage configurations counted and the black dots represent individual flowerbuds (minimum 3). **d)** Expression of pTDM1::TDM1-YFP in metaphase I anthers of Col-0 and tam-2 at 20°C, counterstained with DAPI. For b,d scale bar equals 10µm.

**Figure 8.**
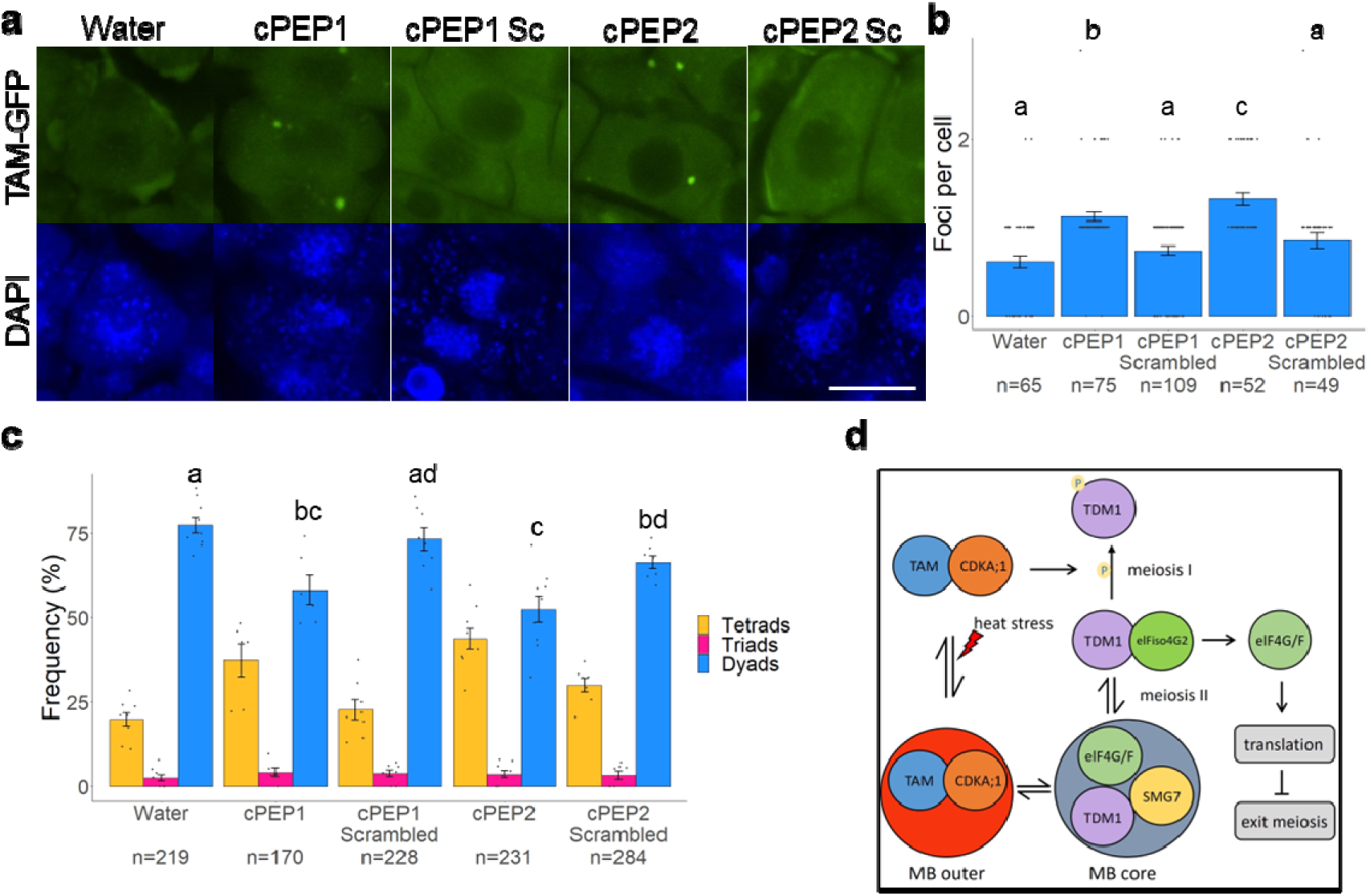
Boosting TAM translation can rescue the HIMR phenotype. Expression of Mt-0’s pTAM::TAM-GFP in prophase meiocytes in the tam-1 background after 24h at 32°C and prior treatment with complementary peptides (cPEP) or a water mock; counterstained with DAPI. Scale bar equals 10µm. **b)** Quantification of the number of TAM-GFP foci for **(a)**. **c)** Frequency of tetrads, triads and dyads for Mt-0 after 24h at 32°C and prior treatment with cPEPs or a water mock. **d)** Schematic model of the meiotic exit pathway under heat stress; explained in text. For (b,c) error bars represent standard errors, n is the number meiocytes or tetrad-stage configurations counted and the black dots represent individual meiocytes or flowerbuds. p-values are based on pairwise Wilcoxon rank-sum tests, Benjamini-Hochberg adjusted for multiple testing.

To assess the biological relevance of these SAAPs, we modelled the complex TAM-CDKA;1 together with TDM1 for Col-0, Mt-0, Nok-3 and the heat-sensitive *tam-1* mutant (Ile283Thr; (Wang et al. 2004)) (Figure 4b). While Col-0 TAM-CDKA;1 is predicted to bind the N-terminus of TDM1 close to its functional, conserved phosphorylation site Thr16 (Cifuentes et al. 2016), Mt-0 TAM, Nok-3 TAM and *tam-1* do not. The changes in the conformations are attributed to TAM sequence variations since the CDKA;1 and TDM1 protein sequences are identical in Col-0, Nok-3 and Mt-0. These data suggest that SAAPs in the natural, heat-sensitive TAM alleles influence the interaction between TAM and TDM1, but we found no indications for a major role of the IDR region.

### High temperature interferes with TAM protein expression

We continued with the functional analysis of five natural TAM alleles: the heat resistant Col-0 allele, and the sensitive Mt-0, Nok-3, Vimmerby and Tac-0 alleles. Transgenic lines expressing pTAM::TAM-GFP of the corresponding five alleles complemented *tam-1* (Figure 2c) showing a similar expression level and foci formation at 20°C (Figure 5a, b; Supp. Movie 1, 2). An end-point analysis after 24h at 32°C, however, showed that the number of foci and overall GFP signal was significantly lower in Mt-0, Nok-3, Vimmerby and Tac-0 compared to Col-0 (Figure 5a, b). Indicating that TAM protein expression is not maintained in heat-sensitive lines.

To study the dynamics of the TAM protein, Col-0 and Mt-0 TAM-GFP were observed during live imaging at different temperatures. At the peak of TAM expression during prophase, the temperature was shifted from 20°C to 32°C. The increase in temperature coincided with an apparent drop in cytoplasmic signal that was initially similar for both alleles (Figure 5c-e). For the Col-0 allele, this was followed by progression into the first meiotic division and a clearing of the TAM-GFP signal by proteasomal degradation (Figure 5c, e; Supp. Movie 3), similar to 20°C conditions (Figure 5d; Supp. Movie 1). For meiocytes expressing the Mt-0 TAM-GFP allele, upon the shift to 32°C, the prophase was prolonged by about 200min compared to Col-0 (Figure 5c, e). This observation corresponds to previous reports on meiotic progression in the *tam-2* mutant (Prusicki et al. 2019). During the extended prophase, the TAM-GFP expression gradually decreased until the first meiotic division was initiated (Figure 5c, e; Supp. Movie 4). These cells did not progress beyond the dyad stage, indicating termination of the meiotic program.

Although TAM condensate formation was shown to be associated with maintaining meiotic progression under heat stress (De Jaeger-Braet et al. 2025), we observed that both heat-sensitive and heat-tolerant TAM variants formed condensates in equal amounts and with similar timing after the temperature shift to 32°C (Figure 5c, f; Supp. Movies 3, 4). Only during the extended prophase in Mt-0 at 32°C did the TAM-GFP levels drop below a certain threshold and did the condensates disappear (Figure 5c, e; Supp. Movies 3, 4). These results suggest that, rather than condensate formation, the TAM protein level is impaired in the natural heat-sensitive lines. Expression of Col-0, Mt-0, Nok-3 and *tam-1* TAM alleles under the 35S promotor in protoplasts heated to 32°C confirmed that all variants are intrinsically capable of forming condensates (Figure 5g). FRAP experiments further revealed that the recovery dynamics between the Col-0 and Mt-0 TAM proteins were similar at 20°C and 32°C (Figure 5h). These results are in line with our *in silico* structural analyses of the natural variants (Figure 4b).

### TAM is not actively degraded at high temperature

The decline in TAM protein level in heat-sensitive accessions is either the result of degradation or their TAM production (see next section) is not sustained. It was previously shown that active degradation of TAM is governed by the proteasome (Cromer et al. 2012). To test if TAM is prematurely degraded during prophase in Mt-0 at 32°C, we treated flowerbuds with the proteasomal inhibitor MG132. The MG132 treatment caused a small increase in TAM levels at 20°C in both Col-0 and Mt-0 (Figure 6a, b). The treatment also induced a small number of dyads, triads and monads (Supp. Figure 9a, b), in line with previous observations that timely TAM degradation is a requirement for meiotic exit (Cromer et al. 2012); altogether showing that our treatment is sufficient to affect TAM degradation. However, fixation-based end-point analysis after 24h at 32°C (Figure 6a, b) and live imaging during the temperature shift (Supp. Figure 9c; Supp. Movie 5) showed that MG132 treatment did not alleviate the reduction of TAM-GFP in Mt-0 at 32°C. This suggests that rather TAM production is not maintained at high temperature during the extended prophase in Mt-0.

### TAM RNA expression is sustained under high temperature conditions

We next investigated how TAM expression is regulated at high temperature. A qPCR analysis on entire inflorescences showed that TAM mRNA expression levels did not decrease upon heat treatment in both heat-sensitive (Mt-0, Vimmerby, Nok-3, Etna-2, Tac-0, tam-1) and heat-tolerant (Col-0) accessions (Figure 6c). On the contrary, we noticed a tendency for a small (<2-fold) overexpression of TAM mRNA in the heat-sensitive accessions and reasoned this might be the consequence of a delay in meiotic cell cycle progression, resulting in relatively more cells residing in prophase. To check if the stage-specific timing of TAM transcription was not affected by heat, we performed mRNA fluorescent in situ hybridisation (mRNA-FISH) to detect the TAM mRNA in whole-mount samples. This experiment was performed in lines expressing pSMG7::SMG7-RFP to account for putative sequestration of the TAM mRNA in MBs at high temperature. In both Col-0 and Mt-0 the TAM mRNA was detected in similar quantities in prophase meiocytes at 20°C and after a 24h at 32°C heat treatment (Figure 6d, e; Supp. Figure 9d). TAM mRNA did not colocalize or associate with SMG7-RFP at either temperature (Figure 6d; Supp. Figure 9e). These results indicate that changes in TAM transcription rate, timing or mRNA degradation at elevated temperature do not explain HIMR in heat-sensitive accessions.

### TAM is required to maintain meiotic translation at high temperature

Because MBs are associated with translational regulation and because active translation is required for meiotic progression, we reasoned that TAM might associate with MBs to regulate meiotic translation during heat stress. To test this we expressed pUBQ10::GFP-RPL18 in Mt-0 and Col-0 ((Thellmann et al. 2020); Supp. Figure 10a) to perform Translating Ribosome Affinity Purification (TRAP). We extracted total RNA and TRAP RNA from whole inflorescences (Supp. Figure 10b) and compared the translation ratio (TRAP RNA/total RNA) within both accessions at 20°C and after 24h at 32°C. We found a lower translation ratio at 32°C in Mt-0 compared to Col-0 for TAM (Figure 7a) and other known meiotic cell cycle regulators (OSD1, TDM1, SDS, PS1, SMG7, CYCB3;1, CDKA;1, CDKD;3), but not for housekeeping genes (RPL18, ACT2, GAPDH, PIN3) (Figure 7a). This shows that heat-sensitive TAM alleles fail to maintain meiotic translation rates at high temperature.

The recruitment of the TDM1 protein into the MB core by SMG7 during the second meiotic division is required for the inhibition of translation that is in turn required for meiotic exit (Cairo et al. 2022, 2025). Since genetic and molecular evidence have shown that the TAM-CDKA;1 complex phosphorylates TDM1 during prophase to prevent a premature meiotic exit (Cifuentes et al. 2016), we wondered if the inhibition of translation under heat in Mt-0 requires TDM1. We therefore performed genetic interaction tests between the *tdm1-3* mutant and the TAM alleles of Col-0, Mt-0 and the *tam-2* mutant. Mutants of *tdm1* are incapable of timely terminating the meiotic cycle, leading to multiple rounds of chromosome segregation and the production of polyad structures with disturbed callose wall formation instead of tetrads at the end of meiosis (Cairo et al. (2022); Figure 7b, c). In agreement with TAM being a negative regulator of TDM1 and *tdm1-3* being epistatic to the *tam-2* mutation (Cifuentes et al. (2016); Figure 7c) we found that the *tdm1-3* mutation supresses Mt-0 dyad production at 32°C (Figure 7c). Interestingly, the 32°C temperature treatment induced dyad and tetrad formation in the *tdm1-3* mutant, independent of the TAM allele (Col-0, Mt-0 or *tam-2*; Figure 7c). Dyads and tetrads are also formed at 32°C in the *smg7-6* and *smg7-6*/*tdm1-3* double mutant (Figure 7c) indicating that heat can terminate meiosis independently of the canonical meiotic exit pathway (Cairo et al. 2022).

TDM1 is present in the cytoplasm of the meiocytes during prophase, and is recruited into the MB core during the second meiotic division to inhibit translation (Supp. Figure 10c; Cairo et al. (2022)). Given that TAM localises around the MB core (Figure 3) and is required to sustain meiotic translation at high temperature (Figure 7a), we asked if TAM prevents translocation of TDM1 to the MB during prophase. Imaging of pTDM1::TDM1-YFP showed that TDM1 condensates are indeed formed during meiosis I in the tam-2 mutant, whereas they only occur during meiosis II in WT plants (Figure 7d; Supp. Figure 10c).

Taken together, these results show that TAM is required to maintain meiotic translation at high temperatures of several cell cycle regulators (including itself), hereby preventing HIMR and 2n gamete formation, through a mechanism that involves TDM1.

### Boosting TAM translation can rescue the HIMR phenotype

If TAM is indeed a regulator of meiotic translation at hight temperature, we considered that artificially boosting TAM translation in Mt-0 at 32°C should suffice to rescue the HIMR phenotype. To do so we employed complementary peptides (cPEPs) that can slightly increase the translation of a selected protein (Ormancey et al. 2023). Their function relies on the presence of TAM mRNA in the treated tissue, a condition that is met in our case (Figure 6c, d). Two synthetic cPEPs that putatively boost TAM protein synthesis were applied to heat treated flower buds expressing Mt-0’s pTAM::TAM-GFP. Treatment with the cPEPs significantly increased the expression of TAM-GFP compared to a water mock (Figure 8a, b). Next, we performed the same treatment on the natural Mt-0 accession. Here the cPEPs treatment lowered the dyad production frequency by about 20% (Figure 8c). These effects were TAM-specific since they did not occur after treatment with cPEPs with the same amino acid composition sequence scrambled in a random order (Figure 8a-c).

## DISCUSSION

Heat stress during sexual reproduction causes a strong reduction in fertility and a range of defects that have been broadly observed among different plants species and crops (Bretagnolle, Thompson, 1995; De Storme and Geelen, 2014). Plants show strong variation in their sensitivity to heat stress, demonstrative for their evolutionary adaptation to the environmental climate conditions. In this study we investigated the natural variation in heat-induced meiotic restitution in *Arabidopsis thaliana*. We identified allelic variants of the core meiotic cyclin TAM/CYCA1;2 responsible for heat-induced dyad formation, the production of diploid male gametes and polyploid offspring. These heat-sensitive alleles occur in late flowering accessions that originate from colder and more variable climates. Late flowering accessions are winter annuals that flower in spring (Shindo et al. 2007; Fournier-Level et al. 2022), whereas early flowering accessions flower in the relatively warmer summer. In a stable climate, summer annuals likely have an adaptive advantage to heat-tolerant meiosis and reproduction (Bac-Molenaar et al. 2015) and thereby maintain ploidy consistency. In the light of the current climate change, winter annuals are more likely to face warm temperatures during their reproductive stage and may suffer more frequently from heat-induced unreduced gamete formation. Polyploidy is considered a driving force for adaptation to harsh and changing environments (Bomblies et al. 2015; Rice et al. 2019) and on an evolutionary scale, the transition from diploid to polyploid species within a clade coincides with periods of global climate change (Van de Peer et al. 2021). The identification of natural, heat-sensitive allelic variants of TAM provides a genetic and molecular mechanism that controls how plants can produce unreduced male gametes and transition into polyploidy.

Translation is dynamic and regulates progression of meiosis in yeast and mammals (Brar et al. 2012; Susor and Kubelka 2017; Sabi and Tuller 2019). In plants, meiotic exit is under TDM1 and SMG7 mediated control of translation (Cairo et al. 2022). TDM1 is suppressed by CDKA;1-TAM phosphorylation at the end of meiosis I, which prevents its premature localisation to MBs and sequestration of translation elongation factors such as elF4isoG2 (working model in Figure 8d). During meiosis II, TDM1 as well as the elF4G/F translation initiation complex are recruited to PBs by SMG7, reducing protein translation and promoting exit of meiosis (Cairo et al. 2022). Our current work shows that TAM is also a component of MBs, and that the protein resides in an interphase surrounding the PB core. Since expression of TDM1 and TAM overlaps during prophase, but TDM1 does not translocate into MBs until TAM is cleared from the cells, we speculate that the MB location of the TAM protein plays a part in preventing TDM1 translocation alongside its phosphorylation function. This observation together with the published genetic interaction between TAM and TDM1 (Bulankova et al. 2010; Cifuentes et al. 2016) suggests that TAM promotes the progression into meiosis II by preventing premature condensation of TDM1 in meiosis I.

In somatic tissue, heat causes a decrease in global translation rates (Yángüez et al. 2013; Merret et al. 2015; Merchante et al. 2017), whereby cell cycle genes are selectively inhibited (Kosmacz et al. 2019; Takahashi et al. 2019), but stress-protection factors are preferentially licensed for translation (Merchante et al. 2017). The untranslated mRNAs trigger the formation of the SG condensates that regulate the storage and degradation of transcripts (Protter and Parker 2016). Our work and that of others have shown that CDKA;1 and TAM translocate to SGs/MBs in response to heat stress (De Jaeger-Braet et al. 2022, 2025). However, TAM condensates form independently of and at temperatures below the formation of canonical SGs. In addition, heat-sensitive and -tolerant TAM alleles are equally able to form condensates, indicating that condensate formation as such is not responsible for heat induced dyad formation. Our results point to a key role of TAM in regulating translation of meiotic cell cycle regulators at high temperature. We propose a model (Figure 8d) whereby heat inhibits meiotic cell cycle gene translation and TAM, together with CDKA;1, is required to move to MBs to regulate this process. This translation regulation is likely indirect and relies on preventing premature TDM1 activation, as evidenced by the epistatic interaction between heat-sensitive TAM alleles and the *tdm1-3* mutation. In accessions with a heat-sensitive TAM allele, TAM translocates to MBs at high temperature but is limited in its functionality. This results in an activation of TDM1-induced inhibition of translation, acting as a negative feedback mechanism whereby cell cycle genes (including TAM) are slowly depleted from the cells until a critical point is reached and the canonical meiotic exit pathway is prematurely activated.

## ACKNOWLEDGMENTS

We would like to thank Ming Yang for providing the *tam-1* seeds and Raphael Mercier for the *tam-2* and *tdm1-3* seeds. We thank Noah Kürtös and Tonni Andersen for providing the UBQ10::GFP-RPL18 material and help with the TRAP protocol. We thank Steven Penfield for the useful discussion on flowering time. We are grateful to Christophe Petit, Ellen Van Gysegem, Patricia Delaere and Jana Pečinková for technical support. This work was supported by the Research Foundation-Flanders (FWO), through a PhD fellowship (Grant No. 11G7421N), and an EMBO scientific exchange grant (No. 11283) to Cédric Schindfessel. Chunlian Jin was supported by CSC grant No. 201606350011. This work was further supported by the Czech Science Foundation (EXPRO grant 23-07969X to K.R.) and we acknowledge the core facility CELLIM funded by MEYS CR (LM2023050 Czech-BioImaging).

## AUTHOR CONTRIBUTIONS

CS designed and performed experiments, analysed and visualised the data, wrote the manuscript and acquired funding. AC designed and performed experiments and wrote the manuscript. PM performed experiments. CJ performed experiments. LLP performed experiments, analysed and visualised the data. PAW designed experiments and reviewed the manuscript. KR designed experiments, reviewed the manuscript and acquired funding. DG designed experiments, wrote the manuscript and acquired funding.

## COMPETING INTERESTS

The authors declare no competing interests

## MATERIAL AND METHODS

### Plant material and growth conditions

For the initial tetrad-stage screen, 212 natural *Arabidopsis* accessions were selected (Lardon et al. 2020). The tam-1 seeds were kindly provided by Ming Yang (Oklahoma State University) and Raphael Mercier (Max Planck Institute for Plant Breeding Research) provided the tam-2 and tdm1-3 lines. Genotyping primers are shown in Supp. Table 5.

Seeds were sterilized using chlorine gas and sown on growth medium (for 1l of medium: 2.154g MS without vitamins, 10g sucrose, 100mg myo-inositol and 0.5g MES, 8g agar. pH adjusted to 5.7. 50mg kanamycin was added for antibiotic selection). Seeds were vernalised for 3 days at 4°C in the dark and moved to 21°C and 16h/8h light/dark for germination. After 5 days, seedlings were transferred to soil substrate (Jiffy) and grown at 20°C under 12h/12h light/dark for 3-4 weeks with regular watering supplemented with liquid fertilizer (Wuxal). Afterwards, plants were grown at 20°C and 16h/8h light/dark without fertilizer to stimulate flowering. Relative humidity was kept at 60%-70%. For experiments after the initial tetrad-stage screen, a 10-week cold treatment at 4°C and 16h/8h light/dark was used for accessions Vimmerby, Nok-3, Tac-0, Ta-0, Etna-2, TOM03, Sorbo, Or-1, Tha-1, before moving them to 20°C to stimulate even flowering.

For temperature experiments, healthy looking, flowering plants with at least 3 siliques on the main inflorescence axis were transferred to a Panasonic MLR-352H-PE versatile environmental test chamber under similar growing conditions, but with different temperatures (20°C, 32°C) for the duration of the treatment (24h).

### Tetrad-stage analysis

Accessions were divided into 54 batches (G1-G54) based on their flowering time under 16h/8h light/dark. Three plants for every accession were kept at 20°C and 10 plants were heat treated for 24h at 32°C. Immediately after the treatment, tetrad-stage flowerbuds of the main inflorescence were dissected and stained with lactopropionic orcein (De Storme and Geelen 2020). For every individual plant the number of tetrad-, triad- and dyad-configurations was recorded. Accessions with at least 3 individual heat treated plants with at least 20 tetrad-stage configuration counts were selected for further analysis (a total of 172 accessions; Supp. File 1). The number of tetrads, triads and dyads were converted to a percentage for every plant and the mean value and standard error were calculated for every accession and treatment. A Kruskal-Wallis rank-sum test on the mean dyad production revealed no batch effect (p=0.32).

### Climate variables analysis

The data on 11 temperature-related climate variables for the earth’s surface from the CHELSEA Bioclim dataset (Karger et al. 2017) were downloaded from https://chelsa-climate.org/. For every *Arabidopsis* accession with a known geographical location (longitude, latitude) the values on these climate variables for their respective location were extracted from the dataset and used for correlation analysis with the dyad production frequency after 24h at 32°C. Data on flowering time were gathered from the Arabidopsis 1001 phenotypes project (Atwell et al. 2010).

### Pollen particle size analysis

A single, freshly opened flower was picked from the main inflorescence and placed in 1ml of Isoton II solution. After vortexing to release the pollen and removal of the flower material the solution was diluted in 10ml of Isoton II and measured using a Beckman Mulitisizer II Coulter counter as previously described (De Storme et al. 2013). For each treatment 3 or 4 individual plants were used that produced 1-3 open flowers on the main inflorescence per day. Analysis based on size thresholds was done using a custom R script. Samples with less than 200 particle counts were removed from the particle size analysis.

### Alexander staining of mature pollen

After dissecting the anthers from nearly opened flowers (before anther dehiscence), viability staining was performed as described by Alexander (1969).

### DAPI staining of mature pollen

To visualise the nuclei in mature pollen grains, open flowers were placed in 200µl of citrate-phosphate buffer at pH 4, with 1% triton X-100 and 1µg/ml DAPI (Vergne et al. 1987). After vortexing and removal of the flower material samples were incubated for 20min in the dark. The solution was centrifuged to pellet the pollen and the pellet was resuspended in 20µl of staining solution before visualisation.

### Plant ploidy analysis

Somatic ploidy determination was done based on the nuclear extraction method described by Galbraith et al. (1983) on fresh leaf material as described by De Storme & Geelen (2011). Propidium iodide stained samples were analysed on a BD FACSVerse flow cytometer.

### Somatic chromosome spreads

DAPI staining of enzyme digested somatic chromosome spreads was done according to Ross et al. (1996) with modifications mentioned in De Storme and Geelen (2020).

### F2 mapping analysis

The F2 generation of a cross between Ler and Mt-0 was heat treated (24h 32°C) during the flowering stage and individual plants were checked for high dyad production at the tetrad-stage. The outcross analysis revealed that the dyad phenotype of Mt-0 is linked to a single recessive locus (Supp. Table 3). Leaf samples of the high dyad producing plants were stored at -80°C before DNA extraction with the Wizard genomic DNA purification kit (Promega). A pooled DNA sample (equal mass) was sent for sequencing on an Illumina NextSeq500 PE150 platform for a total of 36Gb output (predicted coverage: 60x). Adapter and low quality reads were removed with Trimmomatic v0.39 using standard parameters (Bolger et al. 2014). Duplicate reads were removed with FastUniq v1.1 (Xu et al. 2012) and quality was monitored using FastQC v0.11.2 (Andrews et al. 2014). Aligning, mapping and annotation were performed with SHOREmap v3.0, according to Sun and Schneeberger (2015).

### Analysis of SNP frequencies

SNP and indel frequencies for the genomic positions of the TAM locus (chr1: 29081654..29085477) were extracted from the 1001 genomes database (https://tools.1001genomes.org/polymorph/) (Alonso-Blanco et al. 2016), or our own resequencing data in case of Mt-0. Positions of relevant genomic regions and functional protein domains were extracted from EnsemblPlants (https://plants.ensembl.org/index.html) or predicted using the Database of Disordered Protein Predictions (Oates et al. 2013). For the SAAP analysis and *in silico* modelling, TAM sequences were reverse-complemented and translated using the standard amino acid code. All unknown amino acids were marked as “X” and filtered out. Then, accessions were divided according to the detected SAAPs and the dyad formation frequencies were averaged for each group and tested for significant differences withANOVA in R. (R version 4.5.0) The protein structures of the different variants of Arabidopsis TAM-CDKA:1-TDM1 complex were predicted with Alphafold3. The predicted structures were loaded into ChimeraX software (v1.10) for the interface contact analysis and structural visualisation.

### Two-step RT-qPCR

Immediately after the treatment (20°C or 24h at 32°C) all unopened flowerbuds on the main axis were frozen in liquid N_2_ before RNA extraction using the Promega ReliaPrep RNA Tissue Miniprep System. RNA purity and quantity were checked with Nanodrop and 1µg of total RNA was used as starting material for first strand cDNA synthesis using the Promega GoScript Reverse Transcription System with oligo(dT) primers. Two µl of cDNA (1:4 dilution) was added to a 20µl qPCR reaction using the GoTaq qPCR Master Mix (Promega) and ran on a BioRad CFX Opus96 thermocycler. Cycles were as follows: 2min at 95°C, 40x (15s at 95°C and 40s at 60°C). Quality was checked by melting curve analysis and gel electrophoresis of the PCR products. For each accession 3 individual plants were used per treatment (biological replicates) and 3 technical replicates were performed per sample. Data were analysed using the ΔΔCt method as described in Taylor et al. (2019) using the R pcr package (Ahmed and Kim 2018), with EF1αA4 as the reference gene (Ning et al. 2021), and normalized to the 20°C control for each accession. Primers are shown in Supp. Table 5.

### Plasmid construction and plant transformation

Gene cloning was performed with the Multisite-Gateway cloning system (Invitrogen). Primers for cloning are given in Supp. Table 5. For cloning of natural TAM alleles, the promotors were cloned into pDONRP4-P1r and TAM CDSs into pDONR221P1-P2 and combined with a C-terminal GFP moiety into destination vector pK7M34GW. Plasmids were checked through enzyme digestion and sequencing before moving them to Agrobacterium tumefaciens strain GV3130. Floral dip was used to transform all plants. Transformed plants were selected based on antibiotic resistance.

For protoplasts transient expression, the construct pGWB-SMG7-TagRFP was described in Cairo et al. (2022). To generate the construct pGWB441-TAM-YFP, we used the previously cloned TAM CDS and we recombined through LR reaction with the vector pGWB441. To obtain the construct pGWB445-CFP-G3BP-2, we amplified the cDNA of G3BP-2 (AT5G43960) with the primers AtG3BP-like TOPO F and AtG3BP-like stop r. The fragment was introduced into pENTR™/D-TOPO using blunt-end TOPO® Cloning reaction, obtaining the construct pENTR™/D-TOPO-G3BP, which was LR-recombined into the vector pGWB445.

The transgenic lines pTDM1::TDM1-YFP in the tdm1-4 background, pSMG7::SMG7-TagRFP and pRPS5A::TagRFP-RBP47b were previously described (Cairo et al. 2022, 2025).

### Transfection of Arabidopsis mesophyll protoplasts

*Arabidopsis* mesophyll protoplasts were isolated and transfected according to Yoo et al. (2007). Col-0 plants were grown for 4-6 weeks at 22°C under 12h/12h light/dark. 20-30 leaves were cut in fine strips using a razor blade and digested in 15ml enzyme solution (1% cellulase Onozuka R10 (Duchefa) and macerozyme R10 (Duchefa), 0.4M mannitol, 20mM KCl, 20mM MES pH=5.7) for 15min in vacuum followed by 3h in the dark at room temperature. Protoplasts were filtered (70µM) and washed twice in W5 buffer (154mM NaCl, 125mM CaCl_2_, 5mM KCl, 2mM MES pH=5.7) and stored on ice for at least 1h. Then, protoplasts were resuspended in MMg solution (0.4M mannitol, 15mM MgCl_2_, 4mM MES pH=5.7) for a final concentration of 3*10^5^ cells/mL. For transfection, 100µL of protoplasts were gently mixed with 15µg of plasmid DNA and 110µL PEG (4g PEG4000, 2.5mL 0.8M mannitol, 1ml 1M CaCl_2_, 3ml H_2_O), incubated for 10-15min at room temperature before adding 440µL cold W5 buffer. After centrifugation and redissolving the cells in 1mL of W5, the cells were incubated overnight at room temperature before visualisation under the microscope.

For the temperature treatments, 500µL-1mL of transfected protoplasts were placed in a heat block at 32°C (1h) or 39°C (30min) before visualisation. For the cycloheximide treatment, a final concentration of 100µM cycloheximide was added before the temperature treatment. For temperature control on the microscope the Interherence VAHEAT system was used.

### Translating Ribosome Affinity Purification (TRAP)

For the extraction of total and TRAP RNA of *Arabidopsis* inflorescences, the protocol of Thellmann et al. (2020) was adapted. Briefly, about 75 inflorescences were harvested from lines expressing pUBQ10::FLAG-GFP-RPL18 and flash frozen in liquid N_2_. Frozen samples were grinded and homogenized in 5mL polysome extraction buffer (PEB; 5mL 2M Tris-HCl pH=9, 5mL 2M KCl, 5mL 0.25M EGTA pH=9, 1.75mL 2M MgCl_2_, 2.5mL 20% Polyoxyethylene-(10)-tridecyl ether, 2.5mL detergent mix (20% tween 20, 20% triton-x 100, 20% brij 35, 20% igepal in H_2_O), 0.1mL 0.5M dithiothreitol, 0.5mL 0.1M phenylmethylsulfonyl fluoride, 0.1mL 25mg/mL cycloheximide, 0.05 50mg/mL chloramphenicol and H_2_O up to a total volume of 50mL) and centrifuged twice to clear the crude extract. At this point 200µL of extract is aliquoted for total RNA extraction. The rest of the extract is mixed with 60µL of pre-washed anti-GFP magnetic beads for 2h on a stirrer at 4°C. Afterwards, the beads are washed with PEB and 3 times with wash buffer (PEB without detergents) and the beads are collected on a magnetic rack. Beads (TRAP samples) and total RNA samples are then subjected to a TRIzol-Chloroform extraction with overnight precipitation at -20°C for high quality RNA extraction.

### DAPI and SR2200 staining and mRNA-FISH of whole-mount anthers

DAPI and SR2200 staining of male meiocytes in intact anthers was performed as described by Capitao et al. (2021). For consistency and to avoid fading of any fluorescent signal, samples were fixed and prepared for visualisation under a microscope within a single day.

For whole-mount RNA-FISH we adapted a protocol from Huang et al. (2023) for antibody-free labelling of mRNAs using a hybridisation chain reaction (HCR). Custom probes were ordered to detect both the Col-0 and Mt-0 transcript (Molecular Instruments). Briefly, whole inflorescences were fixed in 4% PFA in PBS-T (PBS + 0.1% Triton X-100) for 15min in vacuum and then for another 45min. Subsequently the inflorescences were permeabilised by washing 2x in PBS-T, 2x 10min in methanol, 2x 10min in ethanol, 3x 5min in methanol and 5min each in 75%, 50%, 25% and 0% methanol in PBS-T. At this point the anthers were dissected out of the flowerbuds, followed by a 10min enzyme digestion (0.1g citohelicase, 0.375g sucrose and 0.25g polyvinylpyrrolidone in 25ml water) at 37°C, 3x 5min washing in PBS-T, fixation for 30min in 4% PFA and another 2x 5min washes in PBS-T. Next, anthers were incubated in 200µl pre-heated probe hybridisation buffer (Molecular Instruments) at 37°C and subsequently in 100µl pre-heated probe solution at 37°C overnight (probes were custom designed by Molecular Instruments, sequence information is confidential and proprietary information of Molecular Instruments). Subsequently, the anthers were washed 4x 15min in 500µl pre-heated (37°C) probe wash buffer (Molecular Instruments), 2x 5min in 1ml SSC-T (5x SSC + 0.1% Triton X-100; room temperature) and then in 200µl amplification buffer (Molecular Instruments) at room temperature for 10min under vacuum and then another 20min. Afterwards the anthers were incubated in the hairpin solution (Molecular Instruments) overnight in the dark at room temperature. For the DAPI counterstain, samples were washed 2x 5min in SSC-T, 2x 30min in SSC-T, 1x 5 min in SSC-T, 1x 10 min in PBS-T, incubated 60min in 5µg/ml DAPI and finally washed 3x 5min in PBS-T before mounting in Vectashield antifade medium on a microscopy slide.

### MG132 and peptide treatment of in vitro grown inflorescences

For in vitro cultivation of *Arabidopsis* inflorescences, the protocol of Prusicki et al. (2019) was adapted. After removing the largest flowerbuds, the stem of an inflorescence was cut with a clean razor blade about 0.5cm under the flowerbuds. The inflorescence was positioned in the middle of a well in a 24-well plate filled with 1ml of Apex Culture Medium (ACM; 2.2g/L MS medium, 10 g/L sucrose, 8 g/L agarose, pH 5.8) with vitamins (1000x stock: 10% myo-inositol, 0.1% nicotinic acid, 0.1% pyridoxin hydrochloride, 1% thiamine hydrochloride, 0.2% glycine). The plate was closed and the inflorescences were left to climatise in a growth chamber at 20°C in the light for 4h. Afterwards a volume of 500µL liquid ACM (without agarose) supplemented with 100µM MG132 or an equal volume of DMSO was placed on top of the inflorescences, submerging them completely. Subsequently the plate was moved to a climate chamber set at 20°C or 32°C for a 24h treatment followed by immediate fixation of the samples before analysis.

For the peptide treatments, 100µM of a peptide or an equal volume of water was dissolved in the ACM medium with agarose at the bottom of each well. Cut inflorescences were transplanted in the medium and submerged in 500µL liquid ACM medium supplemented with 100µM peptide or an equal volume of water for 24h at 20°C. The liquid was then removed before a heat treatment of 24h at 32°C, followed by immediate fixation of the samples before analysis. Peptide sequences are given in Supp. Table 5.

### Microscopy

Wide field microscopic images were taken using an Olympus IX81 microscope with an X-Cite 120 LED boost lamp equipped with an Olympus XM10 camera and Olympus Cell M software. Images after Alexander staining were taken on an Olympus BX51 with a ColorView III camera and Cell F software. For confocal microscopy of fixed samples of TAM-GFP, SMG7-RFP and RFP-RBP47b, images were taken on a Nikon A1R HD25 and analysed with Nikon NIS-elements software. The Inverted microscope Zeiss Axio Observer.Z1 with confocal unit LSM 780 was used to image TDM1-YFP in anthers and SMG7-RFP, TAM-YFP and CFP-G3BP-2 in protoplasts. For the anthers, we used the objective C-Apochromat 63x/1.2 W Korr UV-VIR-IR M27. For the protoplasts, we used the objective LCI Plan-Neofluar 63x/1.3 1mm Korr DIC M27. The same system was used for FRAP experiments in protoplasts. In this case, to bleach the regions of interest the 514 nm laser was used at 100% intensity. Signal recovery was measured by time lapse microscopy over the course of 180 seconds.

To perform super-resolution microscopy we used the motorized inverted microscope ZEISS Elyra 7 with lattice illumination pattern for 3D structured illumination (Lattice SIM). We used the objective Plan-Apochromat 63x/1.4 Oil DIC M27 and the 2 x PCO edge sCMOS camera, 1280 x 1280, pixel size 6.5 μm × 6.5 μm. For excitation we used 488 nm and 561nm lasers. The raw images were acquired using a dimension of 1024 x 1024 pixels (pixel size 63 nm) and 9 phases and processed in ZEN Black 3.0 SR (Zeiss), utilizing the 3D SIM2 algorithm, with output sampling = 2 and scaled to raw. The software Imaris 10.2.0 was used for 3D segmentation and volume rendering.

### Live cell imaging

Live cell imaging was performed using Light-sheet microscopy, as previously described (Valuchova et al. 2020). Sepals and petals were carefully removed from approximately 0.4-0.45 mm wide buds and the dissected buds were embedded into a glass capillary (Brand: size 4, blue, ref. number 701910) containing 5MS (5% sucrose in 0.5 Murashige and Skoog Medium including vitamins and MES buffer, Duchefa Biochemie and 1% of low melting agarose, Sigma Aldrich). Samples of TAM-GFP of Col-0 and Mt-0 were developmentally synchronized and placed in one capillary in parallel. Microscopy was performed with a Light-sheet Z.1 microscope (Zeiss). The capillary was placed into the microscope chamber containing liquid 5MS medium and the agarose cylinder including both samples was slightly pushed out of the capillary and submerged into liquid medium in the acquisition chamber of the microscope. Imaging was done using a 20x objective (Detection optics 20x/1.0), single illumination (Illumination Optics 10x/0.2) and one track imaging with a 488 nm laser for GFP in 15 min time increments. An incubation temperature of 21°C or 32°C was used. For the MG132 treatment the experiment began in 5MS medium in 21°C for 30 minutes, then the temperature was increased to 32°C and the medium was simultaneously replaced with MG132 (100µM).

### Statistical analyses

Statistical analyses and methods are detailed in the relevant figure or table captions and the relevant text sections. FIJI (Schindelin et al. 2012) was used for image processing, analysis and quantification.

## Notes

### Competing Interest Statement

The authors have declared no competing interest.

### Summary of Updates

The manuscript was revised adding in silico analyses of the predicted protein structures of natural TAM alleles and the link of amino acid substitutions with the frequency of meiotic restitution. Microscopy of condensate formation with a additional SG marker in protoplasts (RBP47b). Additional natural TAM-GFP alleles were analyzed. Additional live imaging was added, demonstrating the impact of MG132 treatment.

