## Supplemental figures and tables for "Heat stress induces unreduced male gamete formation by targeting meiocyte translation"

### SUPP. FIGURE 1

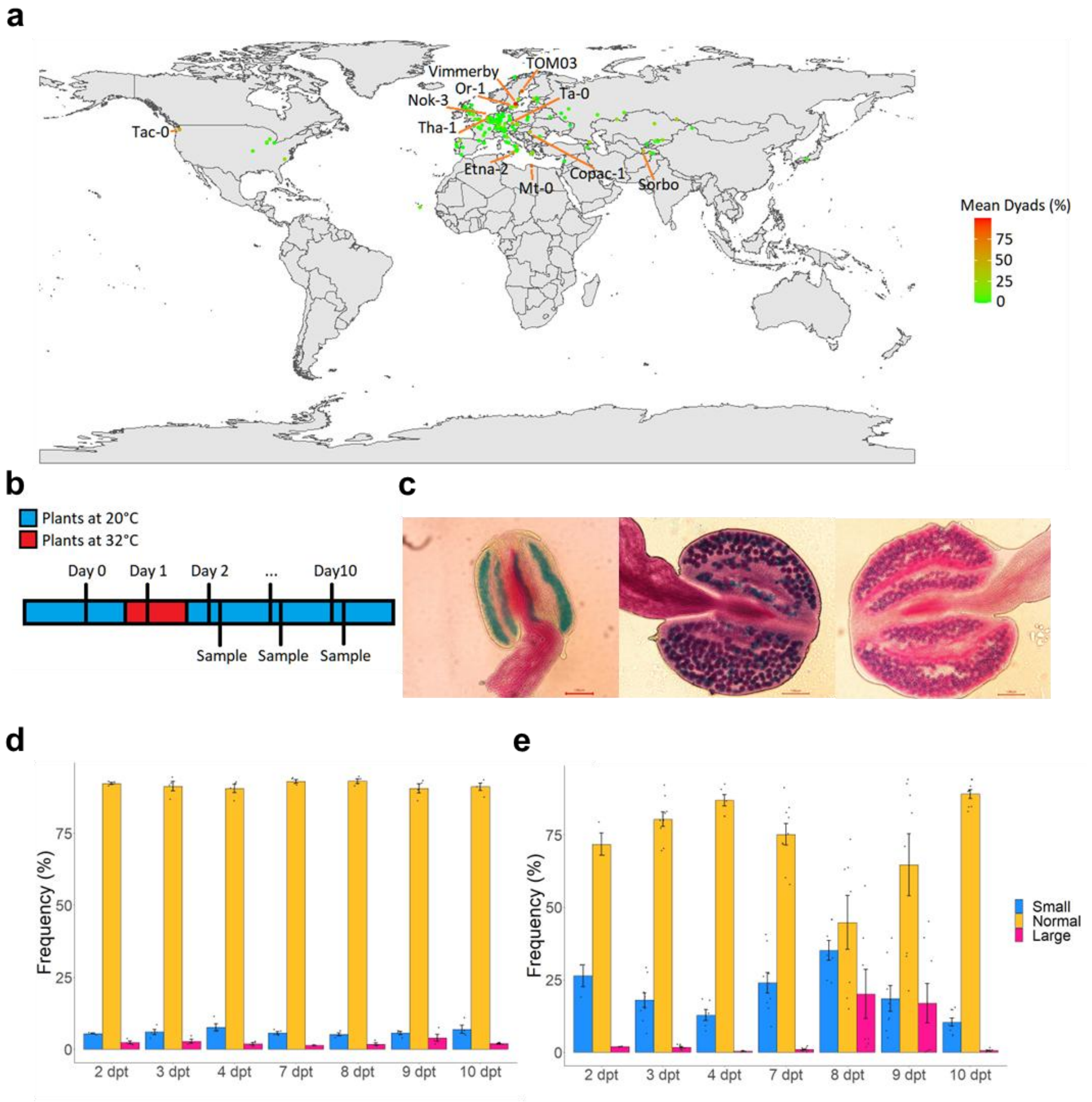

**Supp. Figure 1**

a) Geographical location of the analysed accessions, coloured according to the average amount of dyads produced at the tetrad-stage after 24h at 32°C. The 11 highest dyad producing accessions are denoted with their name. b) Schematic overview of the 24h at 32°C heat treatment, return to the control condition growth rooms and subsequent flower sampling for Coulter Counter analysis. c) Alexander-stained anthers of Mt-0 on days 3 (left), 8 (middle) and 10 (right) after a 24h at 32°C heat treatment. Scale bar = 100µm. d, e) frequency (%) of small (10µm-16µm), normal (16µm-23µm) and large (23µm-30µm) pollen particles analysed through Coulter Counter analysis on 2-10 dpt (days post treatment) for Mt-0 plants kept at 20°C (d) and plants heat treated for 24h at 32°C (e). A minimum of 3 individual plants were analysed for every time point and treatment. Individual flowers are represented by black dots. Error bars represent standard errors.

13

### SUPP. FIGURE 2

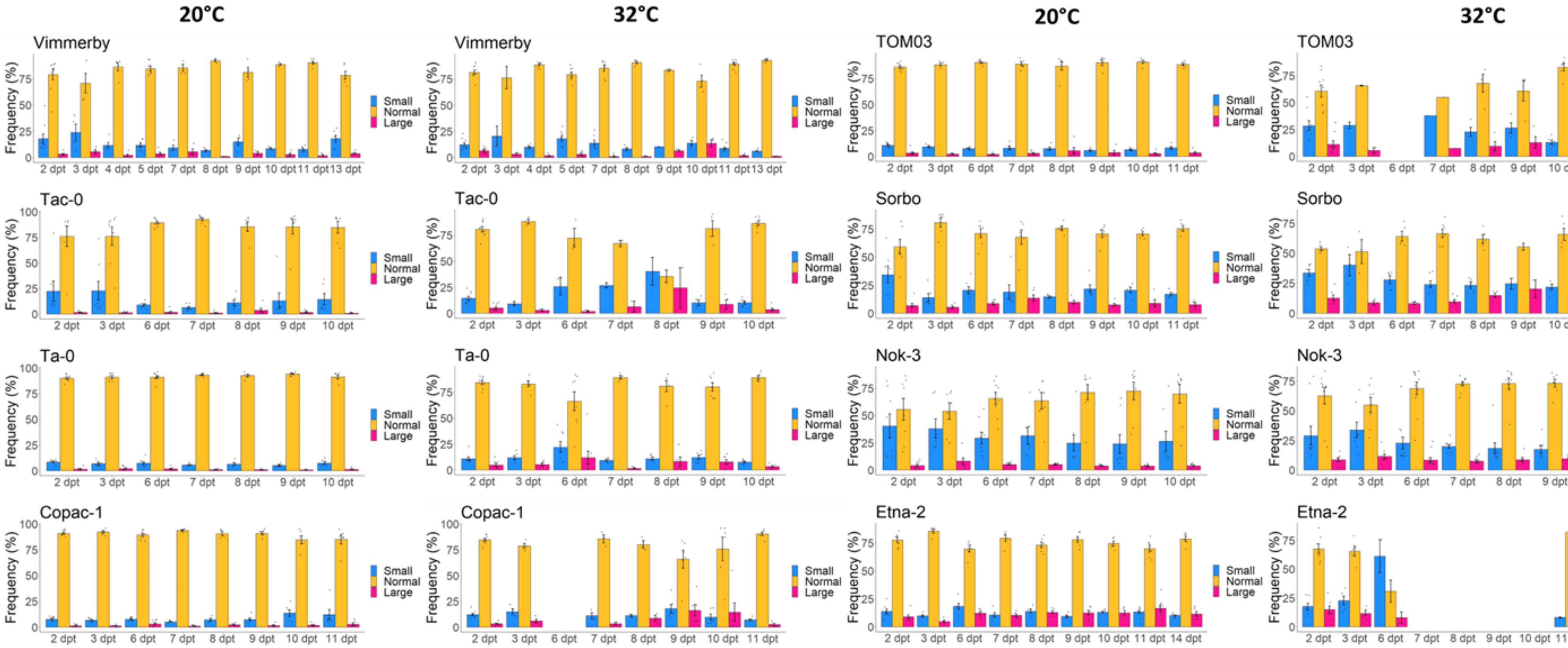

**Supp. Figure 2**

Frequency (%) of small (10µm-16µm), normal (16µm-23µm) and large (23µm-30µm) pollen particles analysed through Coulter Counter analysis for plants kept at control conditions (20°C) and plants heat treated for 24h at 32°C for 8 high dyad after heat producing accessions. A minimum of 3 individual plants were analysed for every time point and treatment. Individual flower measurements are represented by black dots. Error bars represent standard errors. dpt = days after the heat treatment.

20

### SUPP. FIGURE 3

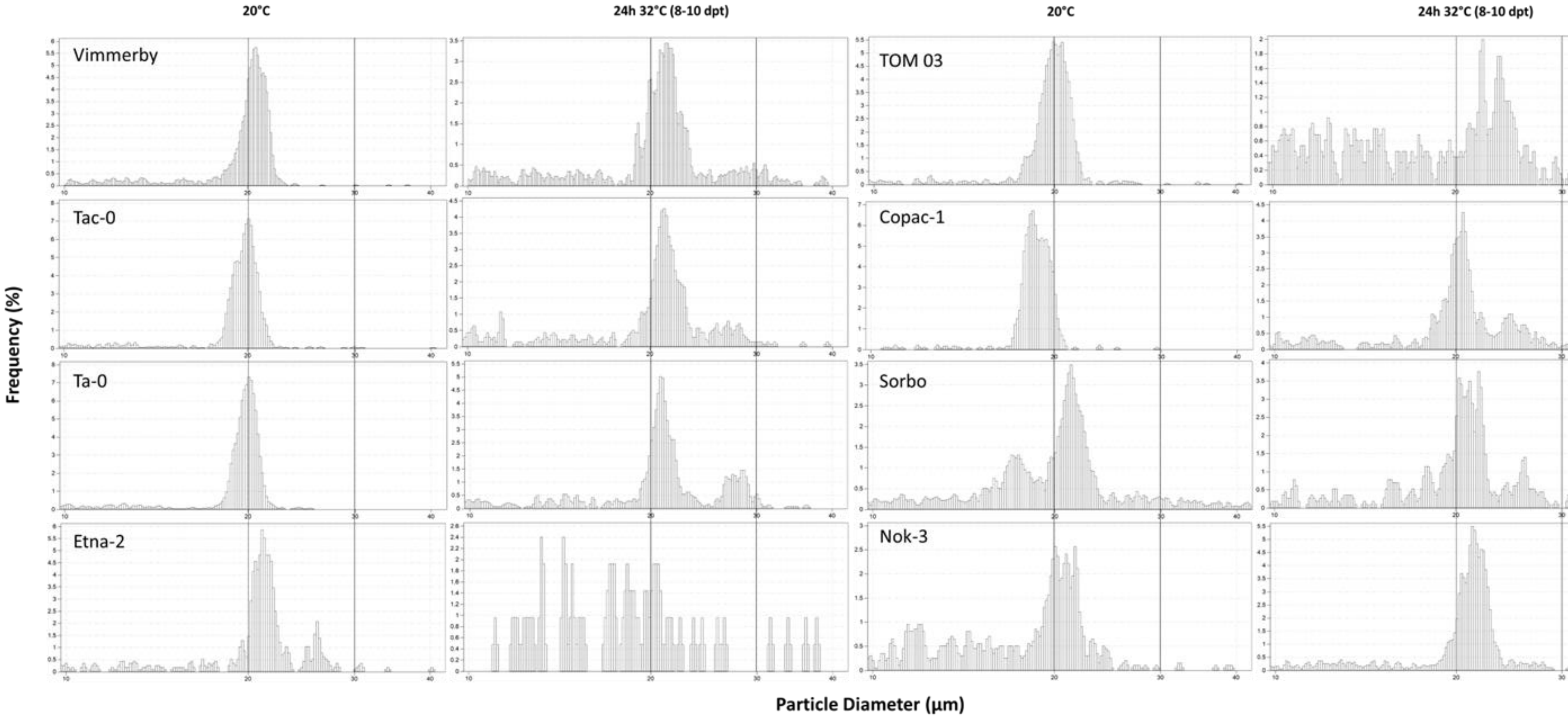

**Supp. Figure 3**

Examples of the pollen particle size distributions of 8 high dyad after heat producing accessions for plants kept at control conditions (20°C) and plants heat treated for 24h at 32°C. The vertical black lines mark the particle sizes of 20µm and 30µm for easy comparison.

### SUPP. FIGURE 4

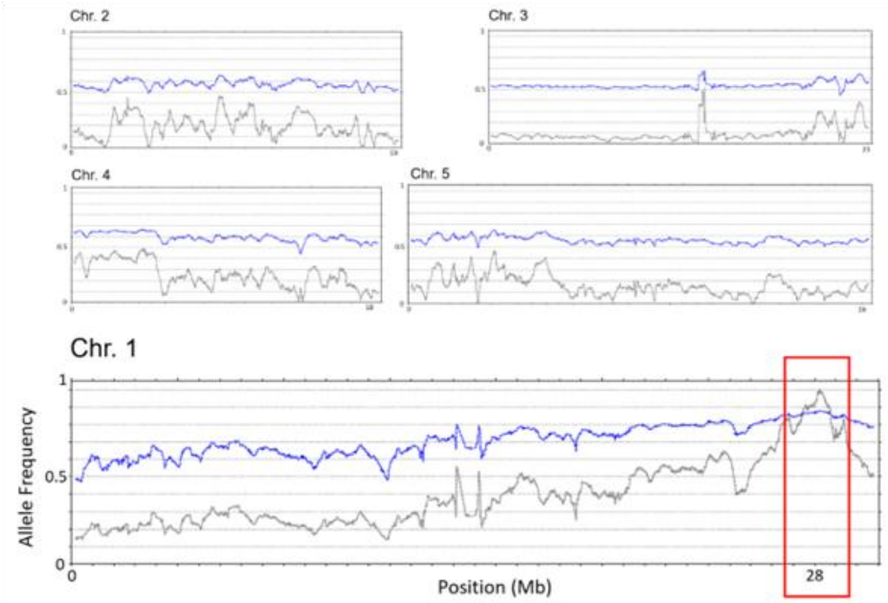

**Supp. Figure 4**

Allele frequency plot of the mapping population for the high dyad phenotype after heat stress of Mt-0xLer. The blue line shows the average allele frequency within a 400kb window (5kb step size), the grey line follows the boost-values (a metric defined by Sun and Schneeberger (2015) to peak sharply at the causal SNP). The red rectangle denotes the predicted 2.4Mbp mapping interval.

#### SUPP. FIGURE 5

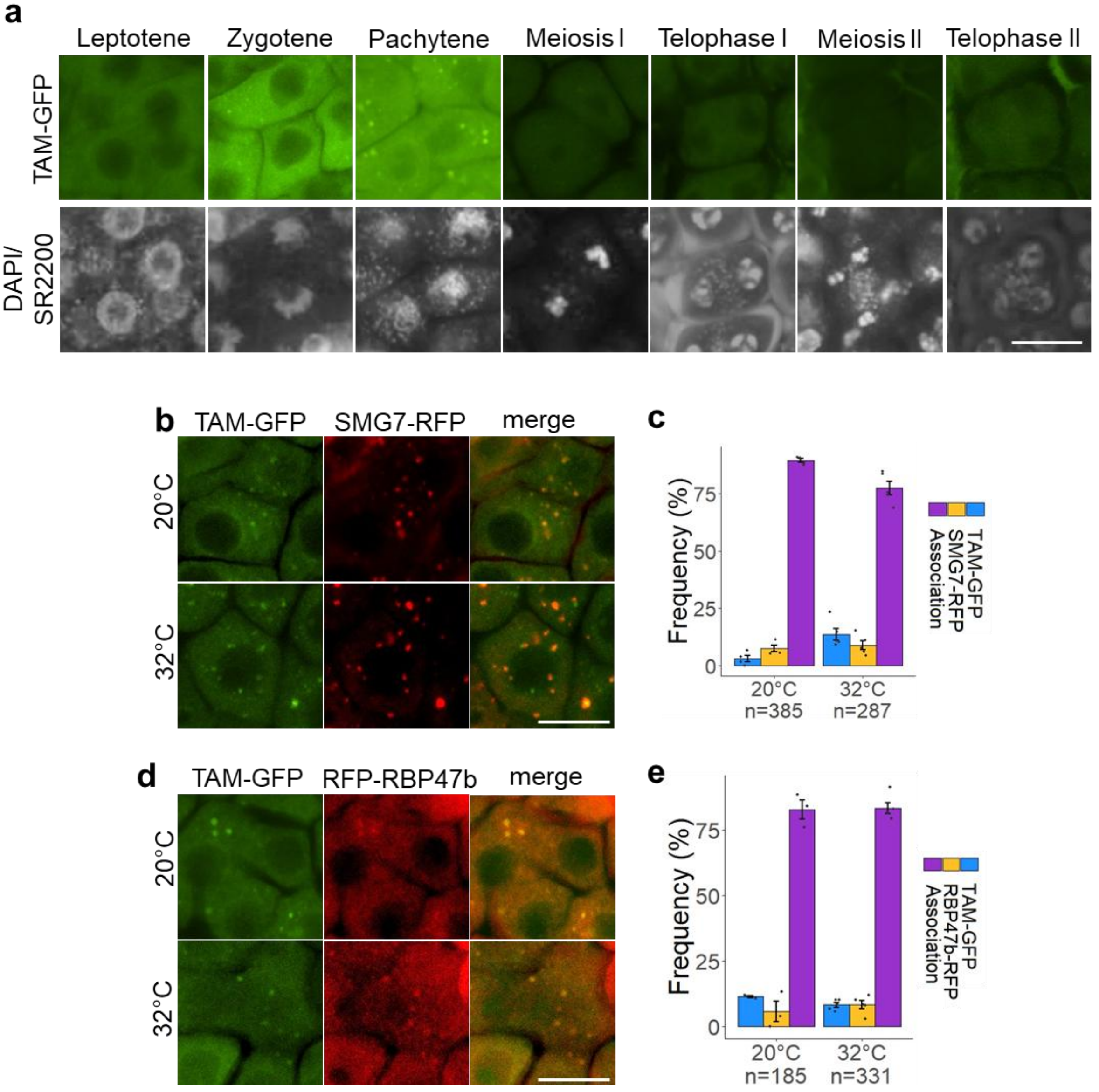


32

**Supp. Figure 5**

**a)** Expression of pTAM::TAM-GFP of Col-0 in the tam-1 background in different meiotic stages; counterstained with DAPI and SR2200. **b,d)** colocalization of pTAM::TAM-GFP and pSMG7::SMG7-tagRFP (b) or TagRFP-RBP47b (d) in prophase meiocytes at 20°C or after 24h at 32°C. Scale bar equals 10µm. **c,e)** Quantification of the association between TAM-GFP and SMG7-RFP (c) or RFP-RBP47b (e) as described for (b,d). Error bars represent standard errors, n is the number of individual foci counted and black dots represent the data for individual anthers (minimum 3). For (a,b,d) scale bar equals 10µm.

**SUPP. FIGURE 6**

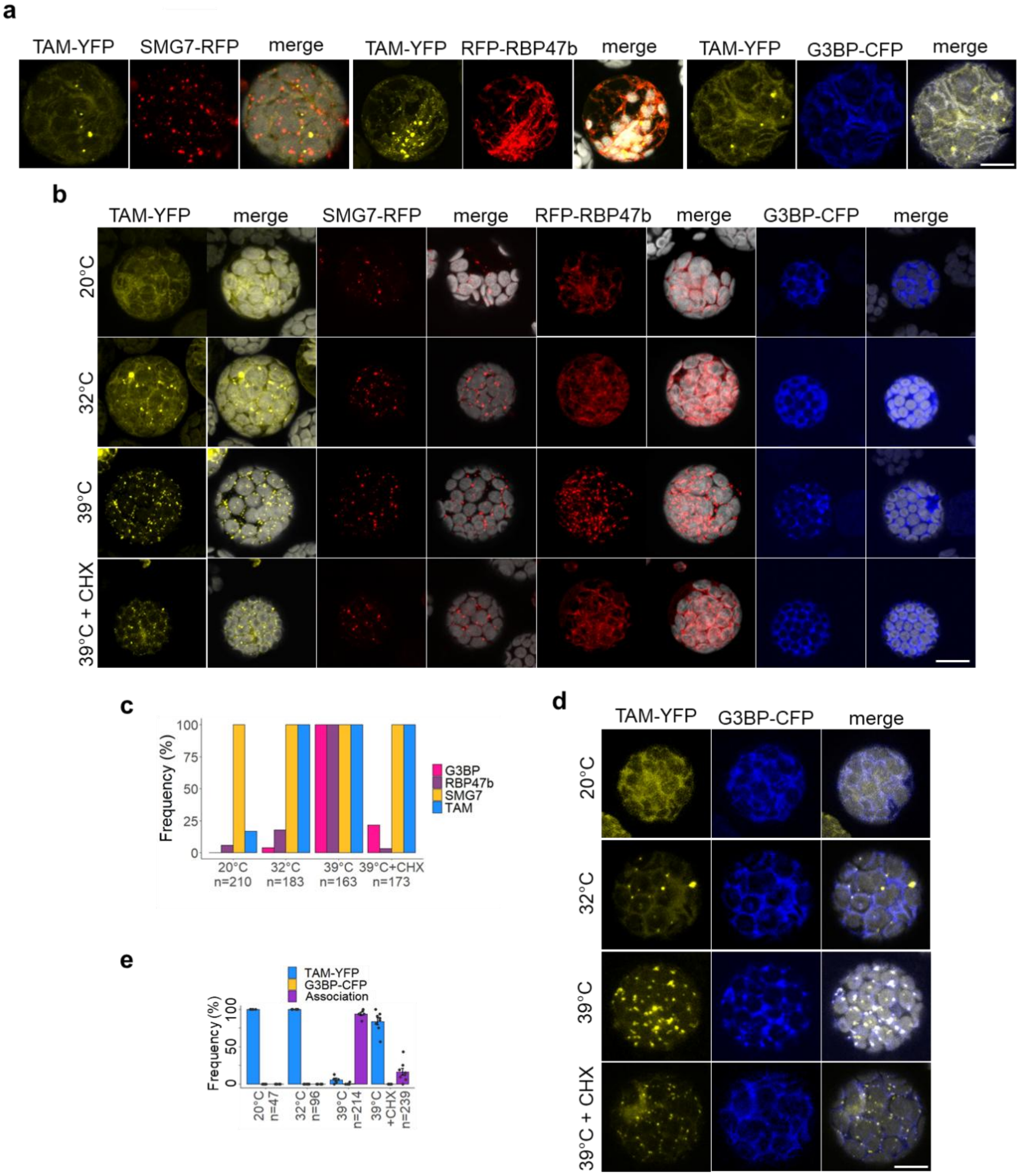

**Supp. Figure 6**

**a)** Co-expression of p35S::TAM-YFP and p35S::SMG7-tagRFP, p35S::tagRFP-RBP47b or p35S::G3BP-CFP in *Arabidopsis* mesophyll protoplasts at 20°C. Examples of cells in which TAM-YFP foci were observed. **b)** Expression of TAM-YFP, SMG7-RFP, tagRFP-RBP47b or G3BP-CFP (not co-expression) in protoplasts at different temperatures and CHX treatment. **c)** Quantification of the number of cells showing TAM, SMG7, RBP47b or G3BP granules in protoplasts at different temperatures and cycloheximide (CHX) treatment for the experiment described in (b). **d)** Expression and colocalization of p35S::TAM-YFP, and p35S::G3BP-CFP in mesophyll protoplasts at various temperatures or treated with 100µM cycloheximide (CHX). **e)** Quantification of the association described in (d). For (c) n is the number of cells counted, for (e) n is the number of foci couted. Error bars represent standard errors and black dots represent individual protoplasts counted (minimum 6). For (a, b, d) scale bar equals 10µm and chloroplast autofluorescence is indicated in grey.

### SUPP. FIGURE 7

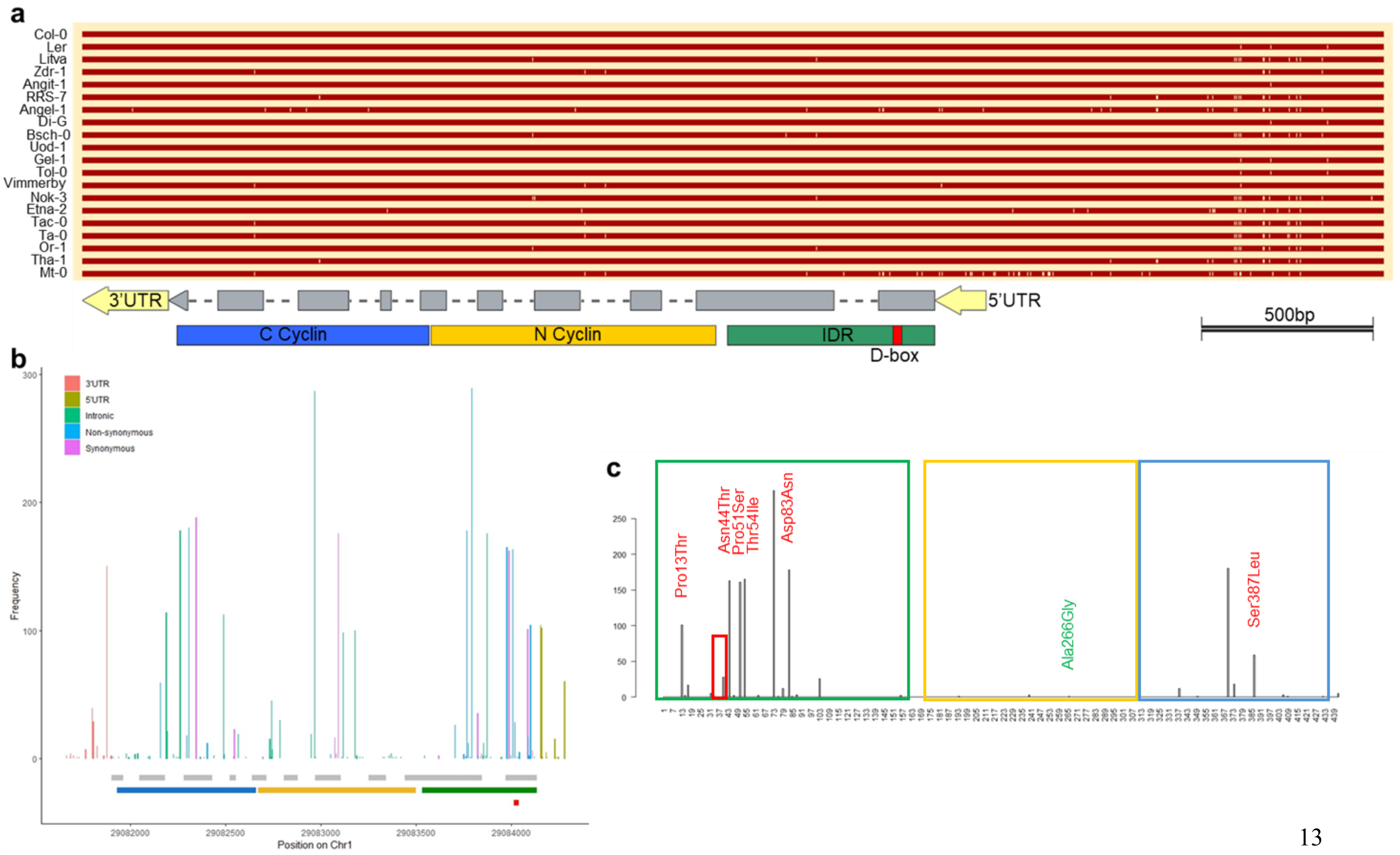

**Supp. Figure 7**

**a)** Alignment of the TAM sequence of the 8 accessions identified to have a heat-sensitive TAM allele (Vimmerby, Nok-3, Etna-2, Tac-0, Ta-0, Or-1, Tha-1, Mt-0), the Col-0 and Ler references and other non-sensitive accessions. The shown region corresponds to the genomic sequence that was cloned for the complementation tests in Figure 2c. White dashes indicate SNPs compared to the Col-0 reference. **b)** SNP frequency across the TAM genomic region, compared to Col-0, across all accessions included in the 1001 genomes project (Alonso-Blanco et al., 2016). **c)** Frequencies of the AA polymorphisms in TAM across all accessions in the 1001 genomes project. Mt-0 non-synonymous SNPs are denoted in Red, Nok-3 non-synonymous SNP is denoted in Green. For (a, b, c) grey boxes represent exons, yellow, blue, green and red boxes represent respectively the N- and C-terminal Cyclin domains, the predicted IDR region and the Destruction box domain (D-box).

#### SUPP. FIGURE 8

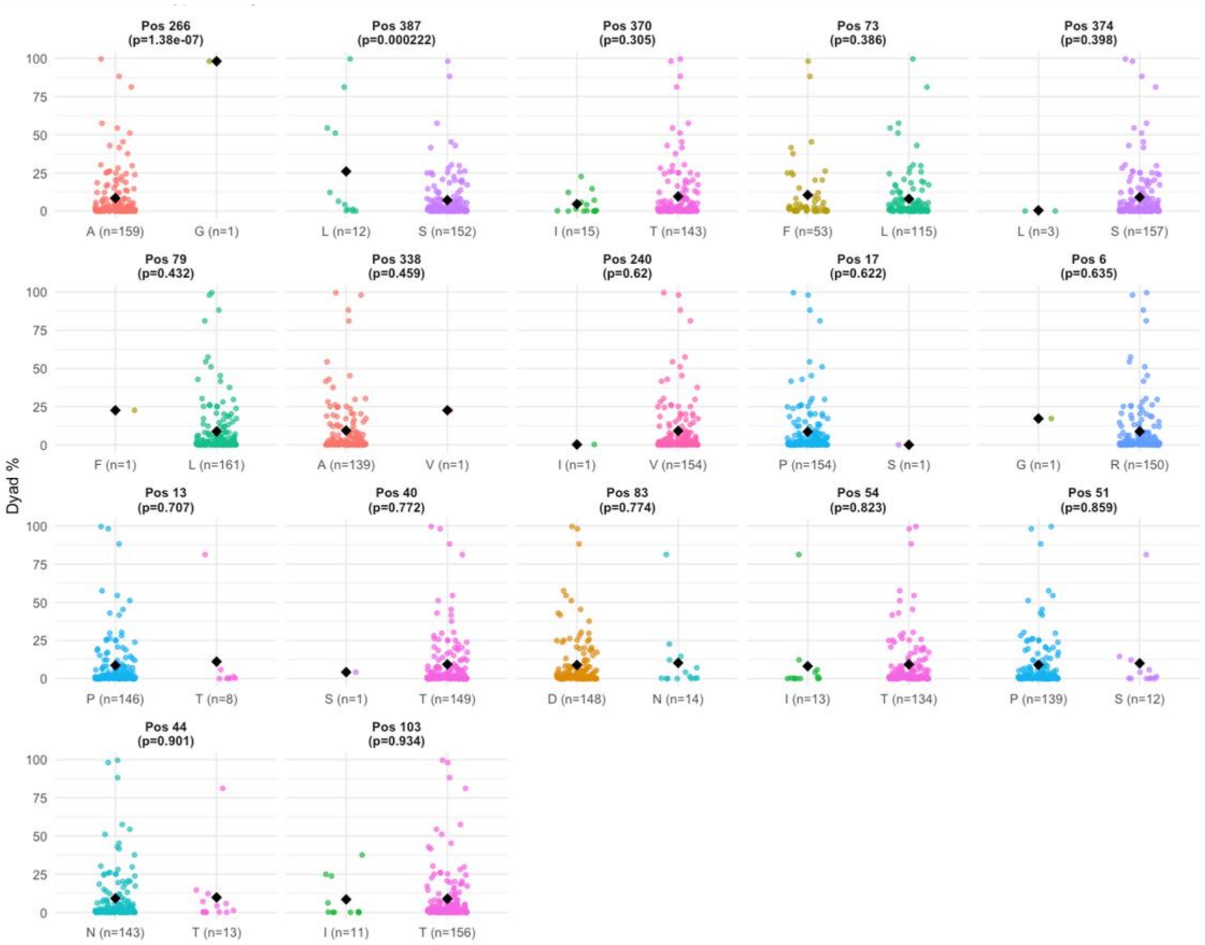

**Supp. Figure 8**

**a)** Comparison of Single Amino Acid Polymorphisms (SAAPs) with dyad frequency after 24h at 32°C (Supp. file 1) for every TAM SAAP in the 172 accessions tested (Sequence data from the 1001 genomes dataset (Alonso-Blanco et al., 2016) and our own sequencing data for Mt-0). p-values based on ANOVA comparing minor and major SAAPs.

**SUPP. FIGURE 9**

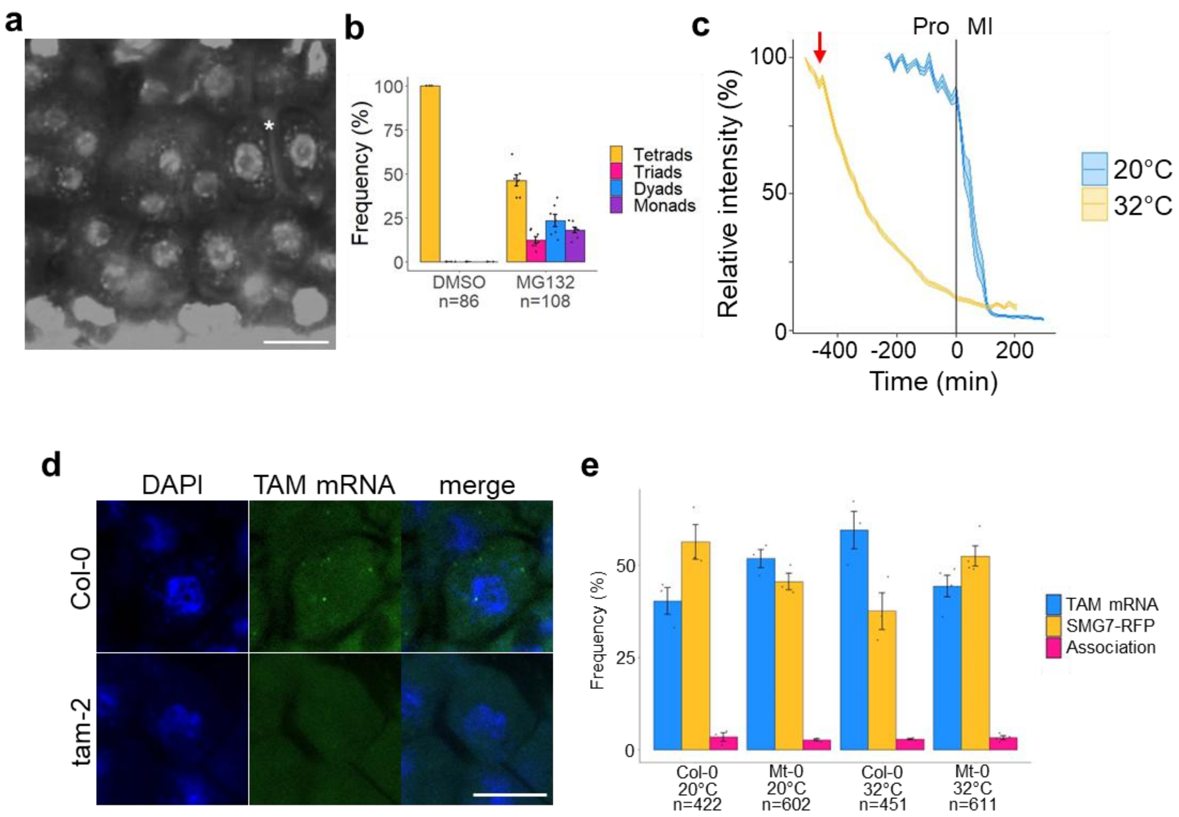

**Supp. Figure 9**

**a)** DAPI and SR2200 stained tetrad-stage anther of Col-0 described in Fig. 6a, treated with MG132 at 20°C, indicating that this treatment sufficiently affected the cells to disturb meiotic progression. Example of a dyad marked with *. **b)** Quantification of the number of tetrads, triads, dyads and monads for (a) and a DMSO mock at 20°C. **c)** Quantification of the fluorescence intensity of TAM-GFP of Mt-0 during a live imaging, time-lapse experiment combining MG132 treatment with a temperature shift to 32°C, cfr. Supp. Movie 5. The red arrow denotes the shift from 20°C to 32°C. The vertical line at time point 0 denotes the entry into meiosis I. Pro = prophase; MI = meiosis I. **d)** Whole-mount RNA-FISH labelling of TAM mRNA, showing that TAM mRNA could be detected in wild type Col-0 plants but not in the tam-2 knock-out mutant. Counterstained with DAPI. **e)** Quantification of TAM mRNA co-localisation with SMG7-RFP, corresponding to Figure 6d. For (a, d) scale bar equals 10µm. For (b) error bars represent standard errors, n equals the number of meiocytes counted and individual anthers are represented by black dots. For (c) n is 18-50 meiocytes across at least 3 anthers and the error ribbon represents standard errors. For (e) error bars represent standard errors, n equals the number of foci counted and black dots represent individual anthers (minimum 3).

**SUPP. FIGURE 10**

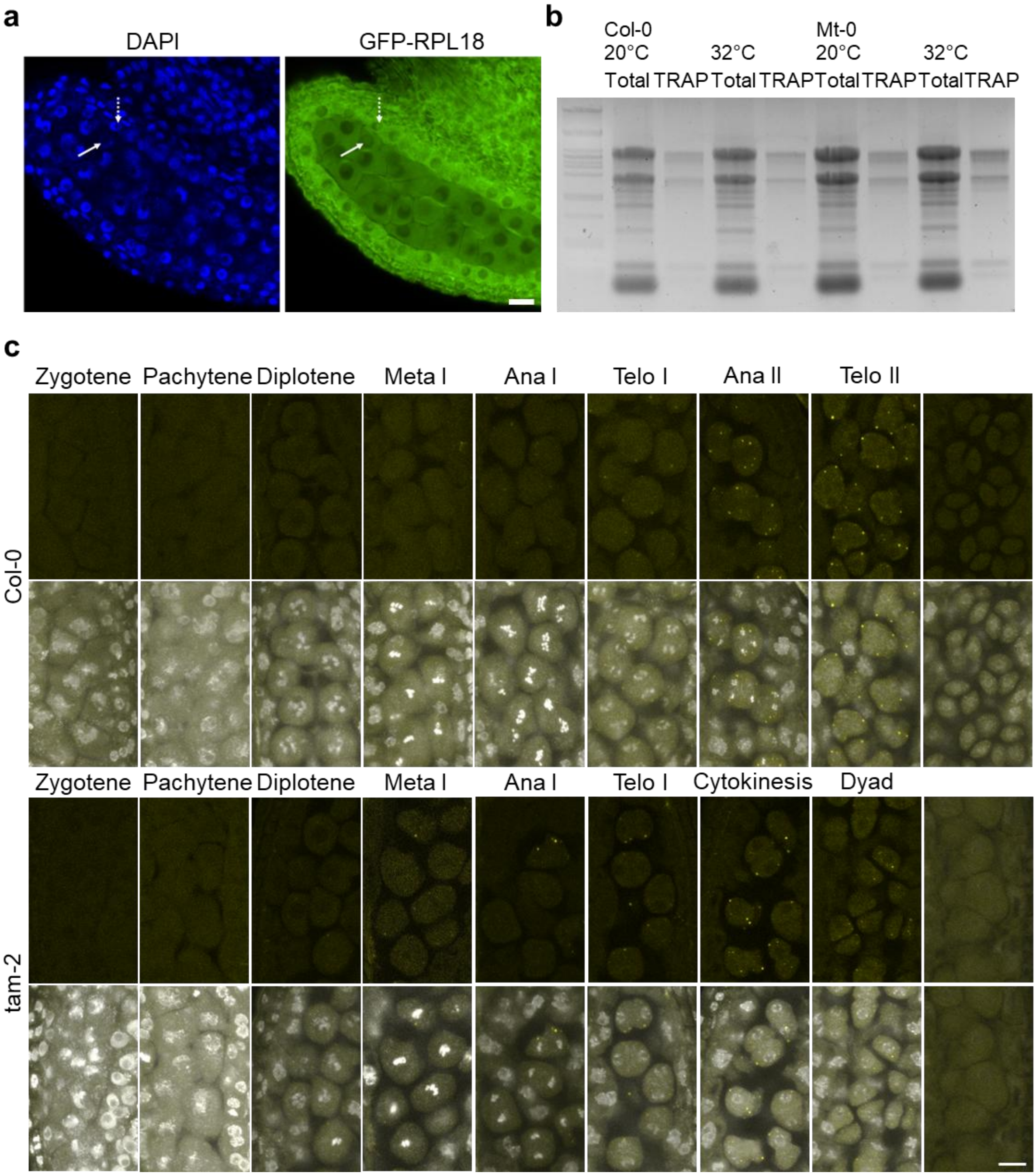

**Supp. Figure 10**

**a)** Expression of pUBQ10::GFP-RPL18 in a prophase *Arabidopsis* anther. The full and dashed white arrow denote the nucleolus region in a meiocyte and tapetum cell, respectively. Counter stained with DAPI. **b)** Agarose (1%) gel showing the RNA fingerprint for total and TRAP RNA extracted from inflorescence of Col-0 or Mt-0 plants expressing pUBQ10::GFP-RPL18. **c)** Expression of pTDM1::TDM1-YFP in anthers of Col-0 and tam-2 at 20°C at different meiotic stages, counterstained with DAPI. For (a, c) scale bar equals 10µm.

| Female Parent | Male Parent | 8dpt | | | 9dpt | | |
|---|---|---|---|---|---|---|---|
| | | 2n | 3n | Germ | 2n | 3n | Germ |
| Mt-0 (24h 32°C; Selfed) | | 2 | 8 | 10/33 | 1 | 4 | 5/50 |
| Mt-0 (24h 32°C) | Mt-0 (20°C) | 17 | 0 | 17/51 | 44 | 0 | 44/51 |
| Mt-0 (20°C) | Mt-0 (24h 32°C) | 77 | 0 | 77/102 | 64 | 15 | 79/102 |

**Supp. Table 1**

The number of diploid (2n) and triploid (3n) plants recorded through flow cytometric analysis coming from a cross whereby Mt-0 after a 24h at 32°C heat treatment was either selfed or used as the male or female parent in a cross with Mt-0 kept at 20°C. Seeds were harvested on siliques that were pollinated 8 or 9 days after the treatment (dpt). 'Germ' denotes the ratio of germinated on sown seeds.

| ShortName | LongName | CorSpear | PVal | PAdjust |
| --- | --- | --- | --- | --- |
| **bio1** | mean annual air temperature (°C) | -0.09 | 0.27 | 0.37 |
| **bio2** | mean diurnal air temperature range (°C) | 0.11 | 0.15 | 0.25 |
| **bio3** | isothermality (°C) | -0.10 | 0.20 | 0.30 |
| **bio4** | temperature seasonality (°C/100) | 0.19 | 0.01 | 0.07 |
| **bio5** | mean daily maximum air temperature of the warmest month (°C) | 0.01 | 0.92 | 0.92 |
| **bio6** | mean daily minimum air temperature of the coldest month (°C) | -0.16 | 0.04 | 0.13 |
| **bio7** | annual range of air temperature (°C) | 0.20 | 0.01 | 0.06 |
| **bio8** | mean daily mean air temperatures of the wettest quarter (°C) | -0.02 | 0.81 | 0.87 |
| **bio9** | mean daily mean air temperatures of the driest quarter (°C) | -0.14 | 0.07 | 0.16 |
| **bio10** | mean daily mean air temperatures of the warmest quarter (°C) | -0.02 | 0.80 | 0.87 |
| **bio11** | mean daily mean air temperatures of the coldest quarter (°C) | -0.14 | 0.06 | 0.16 |
| **FT16** | Flowering time under 16h light | 0.29 | 0.0001 | 0.002 |

**Supp. Table 2**

Climate variables of CHELSEA Bioclim correlated with the mean dyad frequency after 24h at 32°C for all Arabidopsis accessions analysed. Correlations are based on the Spearman correlation and p-values are presented as original and Benjamini-Hochberg adjusted.

| | <5% dyads | >>5% dyads | Total |
|---|---|---|---|
| **Observed** | 730 | 201 | 931 |
| **Expected (1:3)** | 698 | 233 | 931 |

**Supp. Table 3**

The observed and expected number of high (>>5%) and low (<5%) dyad producing plants after 24h at 32°C in a segregating F2 population of Mt-0xLer used for mapping of the Mt-0 locus (considering a single recessive locus).

| | Col-0 DMSO 20°C | Col-0 DMSO 32°C | Col-0 MG132 20°C | Col-0 MG132 32°C | Mt-0 DMSO 20°C | Mt-0 DMSO 32°C | Mt-0 MG132 20°C |
|---|---|---|---|---|---|---|---|
| **Col-0 DMSO 32°C** | 5.37E-01 | - | - | - | - | - | - |
| **Col-0 MG132 20°C** | 7.09E-02 | **9.52E-03** | - | - | - | - | - |
| **Col-0 MG132 32°C** | **2.60E-04** | **6.69E-06** | 7.99E-02 | - | - | - | - |
| **Mt-0 DMSO 20°C** | 1.00E-01 | 2.35E-01 | **8.16E-04** | **8.54E-07** | - | - | - |
| **Mt-0 DMSO 32°C** | **4.58E-33** | **1.13E-34** | **7.43E-43** | **1.23E-50** | **1.82E-18** | - | - |
| **Mt-0 MG132 20°C** | 9.11E-01 | 4.82E-01 | 8.56E-02 | **4.15E-04** | 8.65E-02 | **4.58E-33** | - |
| **Mt-0 MG132 32°C** | **8.01E-25** | **3.31E-25** | **2.88E-33** | **1.71E-40** | **1.01E-13** | 2.09E-01 | **7.35E-25** |

**Supp. Table 4**

Benjamini-Hochberg adjusted p-values for the pairwise comparisons of the quantifications in Figure 6b. Values significant on the 5% significance level are presented in bold.

| Name | Sequence | Purpose |
|---|---|---|
| **TAM_WGS_F3** | CATCACCCATTGCG TGAGAG | Genotyping tam-1 |
| **TAM_WGS_R3** | AATCCTGGTTCATTAAGGCAA G | Genotyping tam-1 |
| **tam-2(SAIL)_LP** | GACTTGATGGATCCACAGC | Genotyping tam-2 |
| **tam-2(SAIL)_RP** | CAGAAATCCTCCACTTGCG | Genotyping tam-2 |
| **tdm1-3_LP** | AGCTTCTGGCTTTTTCGATTC | Genotyping tdm1-3 |
| **tdm1-3_RP** | CCTTACGTCAGAGCCAAACAC | Genotyping tdm1-3 |
| **smg7-6_LP** | CCAGCTCAGACGATTCTCAAC | Genotyping smg7-6 |
| **smg7-6_RP** | TCCATGATTTCCTTGCATCTC | Genotyping smg7-6 |
| **Tdm1-g2** | CACCAGTTCAGGAACCATGACCACAAA | Genotyping tdm1-4 |
| **Tdm1-10** | CAATCTGCATCTGCGTGGTTGTGTA | Genotyping tdm1-4 |
| **Tdm1-5** | TCATTACAACGGCCATATCCTTCAA | Genotyping tdm1-4 |
| **Lbc1** | TGGACCGCTTGCTGCAACTCT | Genotyping tdm1-4 |
| **EF1aA4-qRT-F** | AGGCTGGTATCTCTAAGGATGGTCA | qPCR |
| **EF1aA4-qRT-R** | GGATTTTGTCAGGGTTGTATCCG | qPCR |
| **TAM_qPCR_F** | ACTCTCATCTGATGTGGTTGC | qPCR |
| **TAM_qPCR_R(rvc)** | AAAGCTCTTGCGGTAGTGAC | qPCR |
| **OSD1_qPCR_F** | GTT GCC TTC TTG GTA TCC AAG | qPCR |
| **OSD1_qPCR_R(rvc)** | GTC GAT GAG TTG GGA TCT CAA T | qPCR |
| **TDM1_qPCR_F** | ACT ATG GAA TTG CAG AGC AGC | qPCR |
| **TDM1_qPCR_R(rvc)** | GTC TCG CTC CAA ACC CAA AGC | qPCR |
| **CDKA;1_qPCR_F** | GTA CGA GGA TAC ATG GCG TG | qPCR |
| **CDKA;1_qPCR_R(rvc)** | CTT GAA GTA TTC ATG CTC CAG G | qPCR |
| **SDS_qPCR_F** | ATC CGA CCA AAC TCA ACT CTG | qPCR |
| **SDS_qPCR_R(rvc)** | CTT AAC GCA TTC AGG CAA CTC G | qPCR |
| **CYCB3;1_qPCR_F** | CAC TAC AAT GTC TCC CAG ATG | qPCR |
| **CYCB3;1_qPCR_R(rvc)** | CTT CTC ATA AGT GAC CCT CAG | qPCR |
| **CDKD;3_qPRC_F** | GAC AAG GAT CAA CAA GCA CC | qPCR |
| **CDKD;3_qPCR_R(rvc)** | GAT CGA GTT TCC TCT TCA GAT G | qPCR |
| **SMG7_qPCR_F** | ACC TTG GTA GCT GGT CCT GAG | qPCR |
| **SMG7_qPCR_R(rvc)** | CTG CTG CTA GTG GAG GCA AGA | qPCR |
| **atPS1_qPCR_F** | GAG ATT GAT TCT CCG ACA TCA G | qPCR |
| **atPS1_qPCR_R** | CAT GTG GTG TCT CGC AAA TCA C | qPCR |
| **GADPH-RTF** | ATC AAT GAA GGA CTG GAG AGG T | qPCR |
| **GADPH-RTR** | ATG ACA CCA ACT TCA CAA ACT TGT | qPCR |
| **qPIN3_FW** | GAG GGA GAA GGA AGA AAG GGA AAC | qPCR |
| **qPIN3_RV** | CTT GGC TTG TAA TGT TGG CAT CAG | qPCR |
| **RPL18_qPCR_F** | GAG GTG AAT GCT TAA CCT TTG ACC | qPCR |
| **RPL18_qPCR_R(rvc)** | AGG TCC GAA ATG CTT CAC TGC | qPCR |
| **Actin2-F(qPCR)** | GGC TCC TCT TAA CCC AAA GGC | qPCR |
| **Actin2-R(qPCR)** | CAC ACC ATC ACC AGA ATC CAG C | qPCR |
| **AtG3BP-like TOPO F** | CACCATGGCGACTCCTTATCCTGGAGCG | Cloning |
| **AtG3BP-like stop r** | TTAGCGACCACCACCGCGGTAGTAACC | Cloning |
| **pTAM_gDNAPromotorR4R1_F** | GGGGACAACTTTGTATAGAAAAGTTGTAagagttagagcacgcggtg | Promotor Cloning |
| **pTAM_gDNAPromotorR4R1_R** | GGGGACTGCTTTTTTGTACAAACTTGTgttttctttgattaggggttttctattttatatttattataagaa | Promotor Cloning |
| **pTAM_gDNAPromotorR4R1_R_Mt-0SNPs** | GGGGACTGCTTTTTTGTACAAACTTGTgttttctttgattaggggttttcttttttctatttattataagaa | Promotor Cloning (Mt-0 specific) |
| **pTAM_gDNAPromotorR4R1_F_Nok-3_Tac-0SNPs** | GGGGACAACTTTGTATAGAAAAGTTGTAagggttacagcacgcggtg | Promotor Cloning (Nok-3 and Tac-0 specific) |
| **TAM_gDNA5'UTR_R1R2_F** | GGGGACAAGTTTGTACAAAAAAGCAGGCTTAaaaataattccgcgtttttcattttccgt | Cloning |
| **TAM_gDNAR1R2_noSTOP_R** | GGGGACCACTTTGTACAAGAAAGCTGGGTTgaggaaaagctcttgcggtagtg | Cloning |
| **TAM_gDNA5'UTR_R1R2_F_Mt-0** | GGGGACAAGTTTGTACAAAAAAGCAGGCTTAaaaatgattccgcgtttttcattttccgt | Cloning (Mt-0 specific) |
| **TAM cPEP1** | EDTELLQSDD | Flowerbud treatment |
| **TAM cPEP1 Scrambled** | DEDESLDTQL | Flowerbud treatment |
| **TAM cPEP2** | DNTYLRNELL | Flowerbud treatment |
| **TAM cPEP2 Scrambled** | NYLDERLTNL | Flowerbud treatment |

**Supp. Table 5**
Primers and peptides used in this study.

[_SupplementalFile1.xlsx]
**Supp. File 1**
Information on the 172 accessions retained for analysis in the tetrad-stage screen after 24h at 32°C (HS). Including geographical location (Lat and Long), treatment batch (G1-G54), the number of plants analysed (n) and the mean frequency and standard deviation (SD) of dyad, triad and tetrad production. Ranked according to mean dyad production.

[SuppMovie_1_Col-0TAMGFP_20C.avi]
**Supp. Movie 1**
Representative time-lapse of pTAM::TAM-GFP of Col-0 in the tam-1 background at 20°C from prophase to tetrad stage. Time is indicated in minutes.

SuppMovie_2_Mt-0TAMGFP_20C.avi]
**Supp. Movie 2**
Representative time-lapse of pTAM::TAM-GFP of Mt-0 in the tam-1 background at 20°C from prophase to tetrad stage. Time is indicated in minutes.

[SuppMovie_3_Col-0TAMGFP_TShift.avi]
**Supp. Movie 3**
Time-lapse movie of pTAM::TAM-GFP of Col-0 for the temperature shift experiment described in Figure 5c,e.

[SuppMovie_4_Mt-0TAMGFP_TShift.avi]
**Supp. Movie 4**
Time-lapse movie of pTAM::TAM-GFP of Mt-0 for the temperature shift experiment described in Figure 5c,e.

[SuppMovie_5_Mt-0TAMGFP_TShiftMG132.avi]
**Supp. Movie 5**
Time-lapse movie of pTAM::TAM-GFP of Mt-0 for a temperature shift (20°C-32°C) experiment as described in Figure 5c in combination with an MG132 treatment. t=0 indicates the moment the temperature was increased to 32°C.
